# CTCF-mediated cis-regulatory chromatin insulation enforces a central B-cell tolerance checkpoint

**DOI:** 10.64898/2026.04.12.717890

**Authors:** David Gomes, Eliraz Gitelman, Rena Levin-Klein, Anna Golberg, Ameen Haj-Yahia, Merav Hecht, Bar Avidov, Fernando M. Castellani, Lamia Halaseh, Batia Azria, Arkadiy K. Golov, Natalia T. Freund, Cornelis Murre, Noam Kaplan, Yotam Drier, Yehudit Bergman

**Affiliations:** Department of Developmental Biology and Cancer Research, Institute for Medical Research Israel-Canada, Hebrew University Medical School, Jerusalem, Israel 9112102; Department of Physiology, Biophysics & Systems Biology, Rappaport Faculty of Medicine, Technion – Israel Institute of Technology, Haifa 31096; Department of Computational Medicine, Rappaport Faculty of Medicine, Technion – Israel Institute of Technology, Haifa 31096; The Lautenberg Center for Immunology and Cancer Research, Institute for Medical Research Israel-Canada, Faculty of Medicine, Hebrew University of Jerusalem, Jerusalem, Israel 9112102; Gray Faculty of Medical and Health Sciences, Tel Aviv University, Ramat Aviv, Tel Aviv-Yafo, Israel, 6997801; Department of Molecular Biology, University of California, San Diego, La Jolla, CA, USA, 92093

**Author notes:** Corresponding Authors: Yehudit Bergman, Yotam Drier, and Noam Kaplan E-Mail.

## Abstract

The generation of a diverse and self-tolerant B cell repertoire is essential for adaptive immunity and is achieved through V(D)J recombination. In mice, Igκ is the dominant light chain, whereas Igλ rearrangement typically occurs in response to nonproductive or autoreactive Igκ recombination, a process termed receptor editing. Recombination at the RS element deletes the Igκ constant exon, silencing the locus and enabling Igλ expression. However, the epigenetic regulatory framework that orchestrates and governs receptor editing remains poorly defined. Here, we identify a CTCF-binding insulator element (CBE) within the 3′ Igκ super-enhancer (3′-SEκ) that regulates receptor editing and directs the κ-to-λ switch required for Igλ⁺ B-cell development. Mechanistically, loss of this CBE activates an insulated enhancer within the 3′-SEκ, causing aberrant Vκ rearrangements and altered chromatin interactions through disrupted loop extrusion dynamics. Notably, loss of this CBE in mice leads to increased autoantibody production by ten weeks of age, demonstrating that CBE-mediated chromatin architecture shapes B cell fate by constraining autoreactive potential. Collectively, our findings define a novel CTCF-dependent cis-regulatory insulation checkpoint that connects chromatin loop extrusion to antigen-driven receptor editing, thereby enforcing B-cell tolerance.

## Introduction

The generation of a diverse and self-tolerant B cell repertoire is fundamental to adaptive immunity and relies on the precise orchestration of immunoglobulin (Ig) gene rearrangements^1–4^. Antigen receptor diversity is primarily achieved through V(D)J recombination, wherein variable (V), diversity (D), and joining (J) gene segments are stochastically rearranged to encode functional Ig molecules^5,6^. In developing B lymphocytes, rearrangement of the immunoglobulin kappa (Igκ) locus follows successful assembly of the heavy chain and is subject to stringent spatial and temporal control^1,7^. The murine Igκ locus encompasses over 100 Vκ gene segments alongside four functional Jκ segments, undergoing somatic recombination at the pre-B cell stage^7–12^. This process is tightly regulated by a constellation of cis-regulatory elements that coordinate chromatin accessibility, spatial genome architecture, and recombination potential^13–22^.

Higher-order genome architecture plays an integral role in orchestrating antigen receptor gene assembly^7,23^. Topologically associating domains (TADs) are evolutionarily conserved chromatin domains defined by frequent intra-domain interactions and insulation from adjacent regions^24,25^. Within TADs, finer-scale subdomains known as subTADs often exhibit cell–type–specific interactions that coordinate local gene regulation^24,25^. Both TADs and subTADs are shaped through cohesin-driven loop extrusion and are typically delimited by convergently oriented CTCF Binding Elements (CBEs)^26–34^. The Igκ locus is governed by a multilayered regulatory architecture comprising key enhancer elements, including the intronic enhancer (iEκ)^13,18,19,35,36^, the 3′ enhancer (3′-Eκ)^13–16,19^, and the distal enhancer (dEκ)^14,15,19^. Together with co-enhancers such as HS10^14^ and Dm^17^, these elements form the 3′-Igκ super-enhancer (3′-SEκ)^37^, which regulates chromatin activation and shapes the Vκ gene repertoire. The Igκ locus is organized into CTCF-anchored chromatin loops that define spatial compartments, including subTADs within the Vκ gene array^19,38,39^. Structural boundaries between the Vκ and Jκ domains are reinforced by two prominent CBEs, the Contraction Element for Recombination (CER) and Silencer in the Intervening Sequences (SIS), which insulate the Vκ region from the Jκ recombination centre (RC)^40,41^, thereby limiting proximal Vκ usage and promoting distal rearrangements through long-range chromatin interactions^11,22,42–44^. The CER element enables locus contraction and diffusion-based accessibility required for antigen receptor assembly in murine pre-B cells^40,42^. In the absence of CTCF protein, the Igκ light chain locus similarly exhibits elevated proximal and diminished distal Vκ usage^20^. As B cells progress from early progenitor to precursor B cell stages, the Igκ locus undergoes extensive chromatin remodelling characterized by dynamic shifts in epigenetic marks, transcriptional activation, and the emergence of *de novo* CTCF-mediated chromatin loops that prime the locus for recombination^41,45–48^.

Receptor editing is a vital mechanism of B cell tolerance, allowing autoreactive or nonfunctional B cells to revise their antigen receptor specificity through secondary Vκ–Jκ recombination events or by initiating Igλ light chain rearrangement^49^, which is enabled by the RS element located downstream of the Cκ exon. This process promotes secondary recombination by excising previous Igκ rearrangements^49,50^. Although significant progress has been made in identifying the enhancers and boundary elements that regulate Igκ recombination, the epigenetic mechanisms connecting chromatin architecture to receptor editing and light chain isotype selection remain unknown. In this study, we explore the newly identified function of a previously uncharacterized CBE within the 3′-SEκ, which influences immunoglobulin light chain recombination, isotype balance, receptor editing, and immune tolerance maintenance.

## Results

### CBE is essential for maintaining light chain isotype balance and enabling bidirectional transcriptional insulation

To address whether CBEs located downstream to CER and SIS elements within the 3′-SEκ (Fig. 1a) modulate chromatin architecture or participate directly in Vκ–Jκ recombination, we initially conducted CTCF ChIP–sequencing analysis in IL-7– expanded wild-type murine pre-B cells. We identified two prominent CBEs within the 3’-SEκ region. One element is positioned immediately downstream of the dEκ enhancer (hereafter termed rsCBE), while the other resides within the eighth intron of the ubiquitously expressed *Rpia* gene (hereafter termed rpiaCBE) (Fig. 1a). Both elements display conserved CTCF occupancy, are embedded within a topologically dynamic region of the Igκ locus, and feature binding motifs oriented toward the Vκ domain. Importantly, these elements exhibit conserved CTCF-binding patterns in humans, as evidenced by CTCF-ChIP analysis (Fig. 1b). To dissect their functional roles in Igκ recombination, we employed CRISPR–Cas9-mediated genome editing to generate mice with germline deletion of 571 bp at rsCBE (ΔrsCBE) and 528 bp at rpiaCBE (ΔrpiaCBE). These mouse models allowed us to investigate how the disruption of CTCF-mediated insulation within the 3′-SEκ region influences epigenetic remodelling and recombination dynamics in B cells. First, we assessed whether rsCBE and rpiaCBE modulate the composition of the Igκ repertoire. We performed high-throughput sequencing of RNA from bone marrow-derived pre-B cells isolated from ΔrsCBE and ΔrpiaCBE mice. Comparative repertoire analysis revealed that Vκ gene usage across both mutant lines was essentially indistinguishable from that of wild-type (WT) controls, suggesting that loss of these CTCF-binding elements does not perturb Vκ selection during early B cell development (Extended Data Fig. 1a,b).

**Fig. 1:**
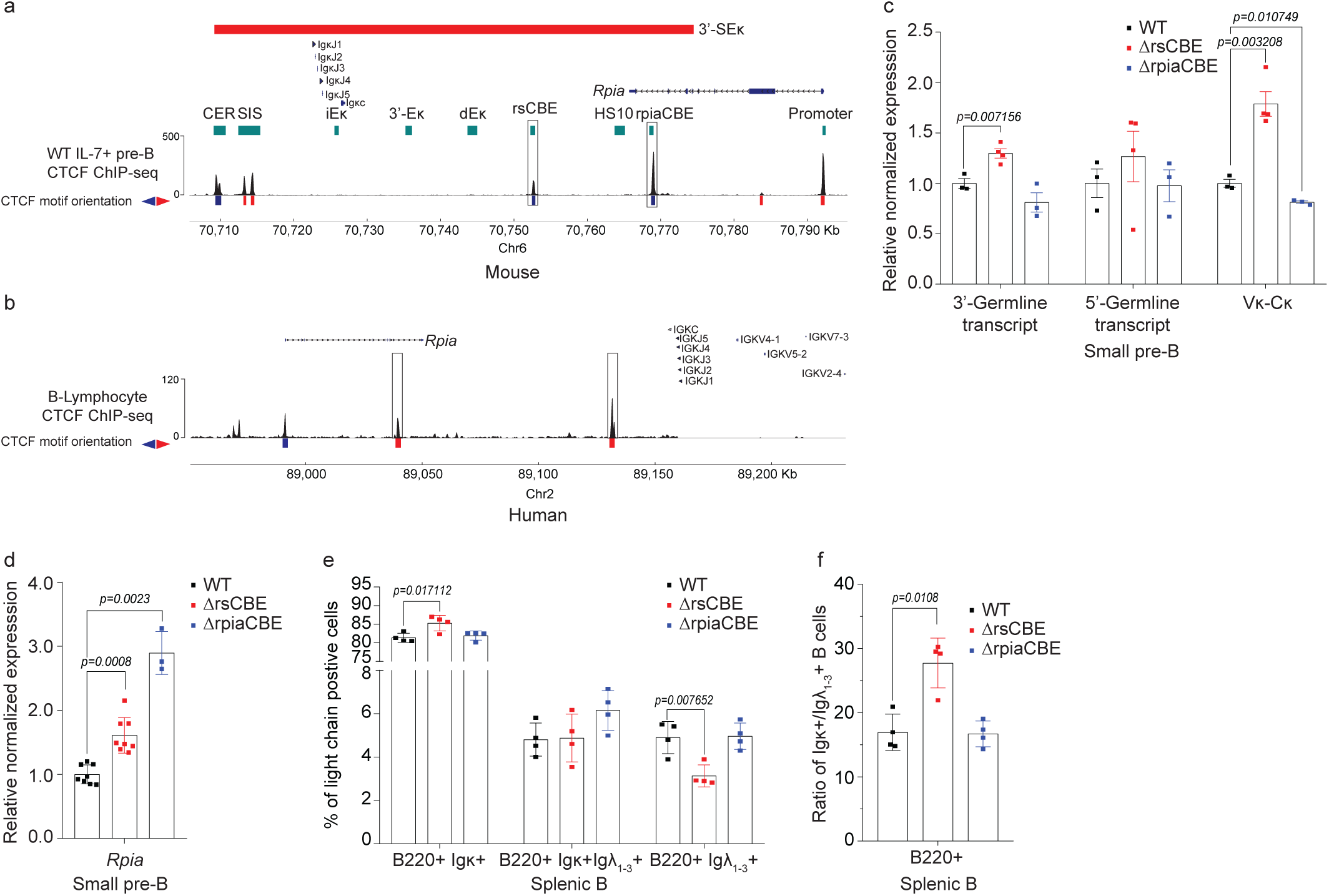
CBE is essential for maintaining light chain isotype balance and enabling bidirectional transcriptional insulation. **a,** Schematic of the 3′-SEκ (red) with CTCF ChIP– seq in IL-7–cultured WT pre-B cells. Regulatory elements and the downstream *Rpia* gene are shown; rsCBE and rpiaCBE (black boxes) were targeted by CRISPR–Cas9 *in vivo*. **b,** Genome browser view of the Igκ locus showing the positions of the RPIA gene and the Jκ region, along with CTCF binding profiles (GSM749762) in EBV-transformed human B lymphocytes. CBEs are highlighted by black boxes. **c,** Quantitative PCR analysis of 3′-GLT, 5′-GLT and total Igκ transcripts in bone marrow–derived small pre-B cells from WT, ΔrsCBE and ΔrpiaCBE mice, normalized to *Ubc* and *Ppia* (n = 3-4). **d,** Quantitative PCR analysis of *Rpia* transcripts in small pre-B cells (n = 3–6). **e,** Igκ and Igλ isotype exclusion in B220⁺ splenic B cells from WT, ΔrsCBE and ΔrpiaCBE mice, expressed as percentages (n = 4). **f,** Ratio of Igκ⁺ to Igλ⁺ B220⁺ splenic cells. Data are presented as mean ± s.d.; P values were calculated using an unpaired t-test.

To assess the impact of CBEs on Vκ–Jκ rearrangement, pre-B cells were isolated from the bone marrow and subjected to quantitative PCR analysis using both genomic DNA and RNA. Deletion of rsCBE led to a pronounced increase in Vκ rearrangement, exceeding 1.8-fold relative to WT controls at the RNA levels (Fig. 1c). In contrast, loss of rpiaCBE resulted in a small yet significant reduction in rearrangement efficiency (Fig. 1c). These findings were also corroborated by increased sterile germline transcripts (GLT) (non-coding transcripts originating from promoters located upstream of Jκ^51,52^) in rsCBE mice (Fig. 1c), and genomic DNA rearrangement closely mirrored RNA expression patterns (Extended Data Fig. 1c). These results collectively indicate that rsCBE, positioned within the 3′-SEκ region, acts as an Igκ chromatin insulator, regulating recombination dynamics.

To assess whether loss of insulation impacts the transcriptional regulation of neighbouring genes, we examined expression of the housekeeping gene *Rpia*^14^, situated approximately 2 kb downstream of the HS10 element (Fig. 1a). Strikingly, disruption of individual CBEs, rsCBE or rpiaCBE, led to a measurable loss of insulation accompanied by an upregulation of Rpia transcripts by ∼1.8-fold and ∼2.8-fold, respectively, in pre-B cells (Fig. 1d). *Rpia* encodes ribose-5-phosphate isomerase A, a pivotal enzyme in the pentose phosphate pathway, and its aberrant expression has been implicated in autophagy^53,54^ and tumorigenesis^55,56^. These data reveal a previously unrecognized role for CBEs within the 3′-SEκ region in safeguarding local transcriptional fidelity, underscoring their importance in preserving the regulatory integrity of the Igκ locus.

Given that approximately 15% of pro-B cells are estimated to initiate Vκ rearrangement^57^, we hypothesized that deletion of rsCBE may induce premature activation of Vκ–Jκ recombination during the pro-B cell stage. In agreement, pro-B cells from ΔrsCBE mice showed a significant over twofold increase in Vκ–Jκ rearrangement compared to WT (Extended Fig. 1d). Notably, rsCBE deletion also resulted in over a twofold upregulation of *Rpia* expression in pro-B cells (Extended Data Fig. 1d). Together, these findings underscore the role of CBEs within the 3′-SEκ region of the Igκ locus as bidirectional transcriptional insulators, functioning to regulate gene expression and recombination events not only in pre-B cells but also at the pro-B cell stage.

Given the altered Igκ expression after CBE deletion, we hypothesized that this may influence the κ⁺/λ⁺ B cell ratio. To investigate this, we first examined the immature B cell compartment in the bone marrow of the ΔrsCBE mouse to ascertain our hypothesis. Interestingly, flow cytometric analysis demonstrated a significant elevation in the proportion of IgM⁺κ⁺ B cells, concomitant with a marked reduction exceeding twofold in the frequency of IgM⁺λ⁺ B cells within the bone marrow of ΔrsCBE mice compared to the control (Extended Data Fig. 1e-g). These findings indicate that rsCBE regulates light chain isotype distribution during early B-cell development, with its deletion skewing the balance toward κ⁺ B cells at the expense of λ⁺ populations in the bone marrow.

We next analyzed splenic B220⁺ cells from the mutant mouse lines to evaluate the role of CBEs in peripheral B-cell populations. Flow cytometry showed that deleting rsCBE caused a significant expansion of the κ⁺ B cell subset, along with more than a twofold reduction in the λ⁺ population (Fig. 1e, Extended Data Fig. 1h,i). As a result, the ratio of splenic Igκ⁺ to Igλ⁺ B cells was significantly increased in rsCBE-deficient mice (Fig. 1f). Importantly, the frequency of Igκ⁺Igλ⁺ double-positive cells remained unchanged between wild-type and rsCBE-deficient groups, indicating that light chain isotype exclusion stays intact. Unlike the rsCBE mutant mice, loss of rpiaCBE alone did not affect light chain isotype distribution (Fig. 1e, Extended Data Fig. 1h,i). Overall, these findings confirm rsCBE as a key regulator of light chain isotype choice in mouse B cells *in vivo*. Thus, our subsequent experiments will focus on the ΔrsCBE mouse, with the ΔrpiaCBE mice serving as a comparison where relevant.

### CBE controls early B cell development

To define the role of CBEs in B cell development, we performed flow cytometric analysis of bone marrow B cell subsets—pro-B, large pre-B, small pre-B, and immature B cells in wild-type and CBE-deficient mice. Deletion of the rsCBE element resulted in a marked reduction in the frequency of small pre-B cells (Fig. 2a, Extended Data Fig. 2a). Conversely, the frequencies of pro-B and large pre-B cell compartments were unaffected significantly, although there was an increasing trend. To determine whether the reduced pre-B cell frequency reflected an absolute numerical loss or a shift in progenitor distributions, we quantified the absolute numbers of each B cell subset isolated from the femur, tibia, and fibula of each mouse. As expected, ΔrsCBE mice exhibited a pronounced decrease in the number of small pre-B cells, whereas pro-B and immature B-cell numbers remained unchanged (Fig. 2b). In contrast, ΔrpiaCBE mice displayed no alterations in B-cell subset distribution (Extended Fig. 2b), suggesting a specific requirement for rsCBE in maintaining early B-cell development.

**Fig. 2:**
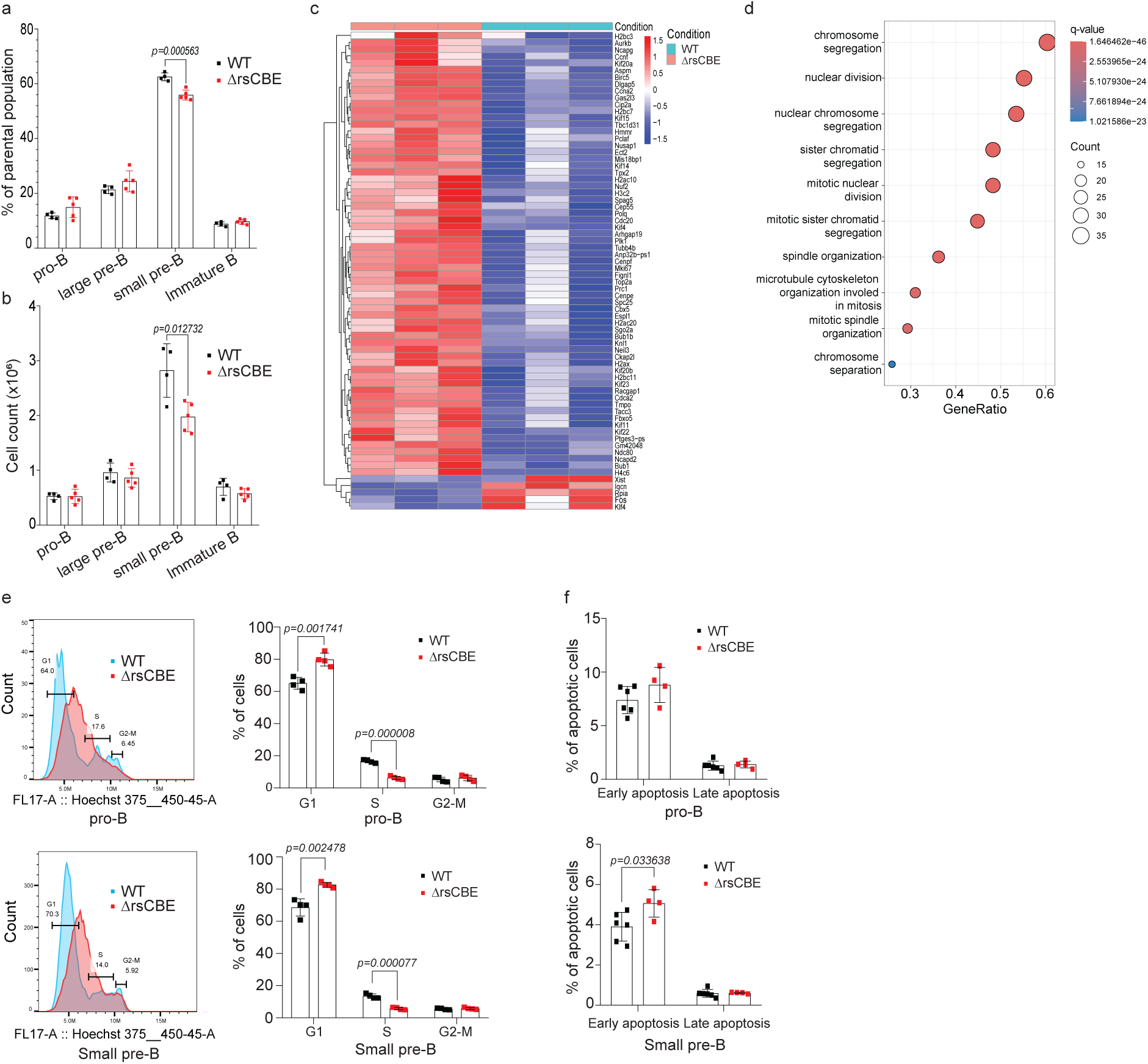
CBE controls early B cell development. **a,** Flow cytometric quantification of bone marrow B cell populations in WT and ΔrsCBE mice, shown as percentages of the parental population; **b,** corresponding cell counts (in millions) (n = 4–5). **c,** Heatmap showing differentially expressed genes (fold change > 1.85, adj. p < 0.1) in small pre-B cells from ΔrsCBE and WT mice. Biological replicates include three WT males and three ΔrsCBE mice (one male and two females). **d,** Gene ontology analysis of downregulated genes in small pre-B cells of ΔrsCBE mice. **e,** Hoechst-based cell-cycle analysis of bone marrow B cell subsets in WT and ΔrsCBE mice; percentages of cells in cycle (G1/S/G2–M DNA content) are shown (n = 4). **f,** Apoptosis in B cell subsets assessed by Annexin V and PI staining; early (Annexin V⁺) and late (Annexin V⁺ PI⁺) apoptotic cells are shown as percentages (n = 4). Data are mean ± s.d.; P values were calculated using an unpaired t-test.

To dissect the developmental impairment in ΔrsCBE mice further, we performed genome-wide transcriptomic profiling of Fluorescence-Activated Cell Sorting (FACS)-purified pre-B cells. RNA-sequencing identified 70 differentially expressed genes, the majority (∼66) of which were significantly downregulated as compared with WT pre-B cells (Fig. 2c). Gene set enrichment analysis of downregulated genes revealed strong enrichment for pathways linked to cell cycle regulation (Fig. 2d). Consistently, cell cycle analysis of bone marrow B-cell subsets by Hoechst staining in WT and ΔrsCBE mice showed a marked accumulation of pro-B, large pre-B, small pre-B, and immature B cells in the G1–S phase, with a smaller fraction in S and G2–M phases, indicating cell-cycle arrest. (Fig. 2e, Extended Data Fig 2c). Given this defect, we next assessed apoptosis using Annexin V staining. In WT and ΔrsCBE mice, pro-B and large pre-B cells exhibited a similar level of apoptosis, while small pre-B and immature B cells showed significantly higher frequencies of Annexin V⁺ early apoptotic cells, indicating elevated cell death (Fig. 2f, Extended Data Fig. 2d). Impaired proliferation combined with increased apoptosis reduces the small pre-B cell compartment in ΔrsCBE mice. This also raises the possibility that the reduction in λ⁺ cells is an indirect consequence of increased cell death among κ-rearranging cells, potentially reflecting enhanced deletion of autoreactive B cells, a possibility that warrants further investigation.

Splenic B cell subsets were analysed by flow cytometry in ΔrsCBE and ΔrpiaCBE mice, revealing no significant differences in Marginal Zone B (MZB), Follicular B (FoB), or Transitional 1 B (T1B) cells (Extended Data Fig. 2e). Altogether, these results identify rsCBE as a pivotal cis-regulatory element that sustains proliferation and developmental progression of early B cell precursors. Its loss disrupts cell-cycle dynamics and diminishes pre-B cell output, a phenotype not observed upon rpiaCBE deletion, highlighting a unique role for rsCBE in safeguarding pre-B cell expansion.

### Co-ordinated molecular epigenetic mechanisms orchestrate Igκ expression

Previous studies have demonstrated a strong correlation between chromatin accessibility and the efficiency of Vκ–Jκ rearrangement^58–62^. To explore changes in the epigenetic landscape across the Igκ locus, we performed ATAC-seq on *ex vivo*– purified pre-B cells. Our analysis specifically examined whether deletion of rsCBE, and the consequent loss of insulation, alters chromatin accessibility both upstream and downstream of the deleted regions. Given that the pre-B cells were derived from Rag-proficient mice lacking an intact germline configuration, our investigation concentrated mainly on the 3′-SEκ region of the Igκ locus. Loss of rsCBE-mediated insulation resulted in increased chromatin accessibility downstream of the deletion site, unveiling a previously insulated regulatory element, termed rsEκ (Fig. 3a). Notably, rsEκ displayed significantly greater accessibility in ΔrsCBE mice compared to wild-type controls and was markedly more accessible than the neighbouring HS10 co-enhancer element (Fig. 3b). In contrast, deletion of rpiaCBE induced a modest, non-significant decrease in accessibility across the three canonical enhancers (Fig. 3a,b). These results reveal a complex interplay between the rsCBE and other regulatory elements in shaping the Igκ chromatin accessibility landscape.

**Fig. 3:**
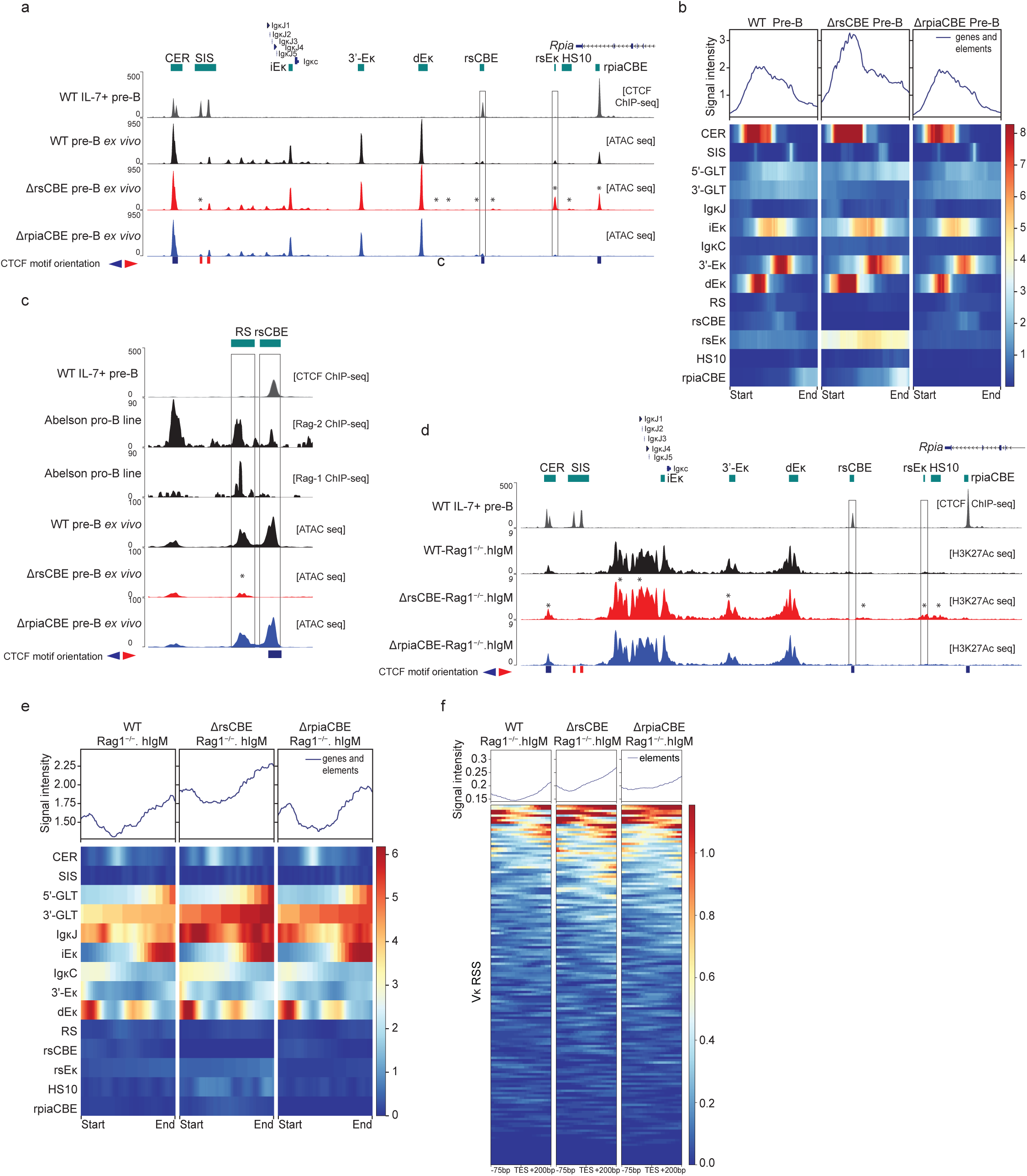
Co-ordinated molecular epigenetic mechanisms orchestrate Igκ expression. **a,** Genome browser tracks across the Igκ locus showing CTCF ChIP–seq from IL-7–cultured WT pre-B cells and ATAC–seq signal from WT-Rag1⁻/⁻.hIgM, ΔrsCBE-Rag1⁻/⁻.hIgM, and ΔrpiaCBE-Rag1⁻/⁻.hIgM pre-B cells. Peaks were called with MACS2 (FDR < 0.05); differential peaks were tested using edgeR (P < 0.05, fold change > 1.5); asterisks denote significance (n = 2). **b,** Heat map of ATAC–seq signal intensity across regulatory elements within the 3′-RR of the Igκ locus. **c,** Browser view of the 23-RSS within the RS region showing CTCF, Rag1 (GSM4176366) and Rag2 (GSM4176369) ChIP–seq, together with ATAC–seq signal in WT-Rag1⁻/⁻.hIgM, ΔrsCBE-Rag1⁻/⁻.hIgM, and ΔrpiaCBE-Rag1⁻/⁻.hIgM pre-B cells; significant differences are indicated by asterisks. **d,** Genome browser tracks showing CTCF ChIP–seq from WT pre-B cells and H3K27ac ChIP–seq from WT-Rag1⁻/⁻.hIgM, ΔrsCBE-Rag1⁻/⁻.hIgM, and ΔrpiaCBE-Rag1⁻/⁻.hIgM pre-B cells. Peaks were identified with csaw; differential peaks were assessed by edgeR (FDR < 0.05, fold change > 1.5; n = 2). **e,** Heat map of H3K27ac enrichment across regulatory elements within the 3′-RR. **f,** Heat map of H3K27ac enrichment at 12-RSS across Vκ gene segments in WT-Rag1⁻/⁻.hIgM, ΔrsCBE-Rag1⁻/⁻.hIgM, and ΔrpiaCBE-Rag1⁻/⁻.hIgM pre-B cells, quantified within a 275-bp window (75 bp upstream and 200 bp downstream of transcription end site (TES) of Vκ genes). Data are based on the mean of two biological replicates from female mice.

CTCF-binding elements are established regulators of RAG substrate accessibility^21^. Given that the rsCBE, a CTCF-binding site, lies ∼160 bp downstream of the RS element, we assessed chromatin accessibility at the RS RSS. Of note, deletion of rsCBE caused a pronounced decrease in accessibility at the Rag1 binding site within the RS element (Fig. 3c). These findings reveal that the structural integrity of rsCBE is crucial for maintaining recombination signal sequences (RSS) accessibility at the RS locus, and its disruption impairs Vκ–RS recombination efficiency (see below).

To assess whether alterations in chromatin accessibility correlate with changes in active enhancer marks, we evaluated H3K27ac enrichment. Wild-type and CBE-deleted mutant mice were bred onto a Rag1-deficient background and crossed with human IgM (hIgM) transgenic mice. This strategy ensured that the Igκ locus remained in the germline configuration at the pre-B cell stage. Bone marrow-derived CD19⁺ cells, composed of over 95% pre-B cell purity, were isolated from WT-Rag1⁻/⁻.hIgM and mutant-Rag1⁻/⁻.hIgM mice and subjected to H3K27 acetylation ChIP sequencing.

Since increased acetylation has been correlated to greater recombination efficiency^63–65^, we first evaluated its enrichment at the 3′-SEκ. Notably, within the 3’-SEκ, the rsEκ and HS10 elements showed hyperacetylation only in ΔrsCBE-Rag1⁻/⁻.hIgM mice (Fig. 3d,e). In ΔrsCBE-Rag1⁻/⁻.hIgM cells, elevated H3K27 acetylation extended to the CER, 5′-GLT and 3′-GLT promoters, RC, and the canonical 3′-Eκ regions. By contrast, ΔrpiaCBE-Rag1⁻/⁻.hIgM pre-B cells showed no significant changes in H3K27ac across these regulatory elements within the 3′-SEκ (Fig. 3d,e). H3K27ac enrichment was also markedly increased across the Ig Vκ gene segments in ΔrsCBE-Rag1⁻/⁻.hIgM pre-B cells as compared to WT (Extended Data Fig. 3a). We also assessed the H3K27ac levels at RSS in the Vκ region. We found increased acetylation at the 12-RSS across the Ig Vκ gene segments in ΔrsCBE-Rag1⁻/⁻.hIgM pre-B cells (Fig. 3f). We observed no significant changes in H3K27 acetylation at the Ig Vκ gene segments or their RSS in ΔrpiaCBE-Rag1⁻/⁻.hIgM pre-B cells compared with WT. These results identify rsEκ as a previously insulated putative shadow enhancer that gains chromatin accessibility and active histone modifications upon loss of rsCBE-mediated insulation. Together, our findings reveal that disruption of a specific CTCF-mediated insulation within the 3’-SEκ reshapes chromatin architecture, influencing Igκ locus regulation.

### CBE controls the intricate Igκ super-enhancer architecture through chromatin insulation

Within the Igκ locus, the 3′-SEκ orchestrates chromatin interactions necessary for proper light chain recombination and B-cell development^13–19^. CBEs play a pivotal role in constraining these interactions through mechanisms such as chromatin insulation^11,20,22,41,42,44^, yet how they influence the internal configuration of super-enhancer domains remains incompletely understood. To investigate the chromosome topology of the 3′-SEκ architecture and its relationship to Igκ recombination, we performed Micro-C-TALE in WT-Rag1⁻/⁻.hIgM and ΔrsCBE-Rag1⁻/⁻.hIgM mice. Micro-C-TALE is a BAC-based targeted chromatin conformation capture method, which utilizes micrococcal nuclease digestion of cross-linked chromatin to enable high-resolution profiling of chromatin folding at target regions. We applied this approach to *ex vivo* purified CD19⁺ pre-B cells to map chromatin organization across the Igκ locus with fine structural detail. Micro-C-TALE yielded approximately 7-12 million high-quality paired-end reads per sample, producing high-resolution contact maps across biological replicates from WT-Rag1⁻/⁻.hIgM and ΔrsCBE-Rag1⁻/⁻.hIgM pre-B cells. While chromatin contact frequencies across the Igκ variable region were broadly similar in both genotypes (Extended Data Fig. 4a), local short-range interaction differences emerged upon detailed inspection of the contact maps at the 3′ regulatory region (3′-RR) of the Igκ locus (Fig. 4a). Notably, we observed genotype-specific contact frequency at key regulatory nodes involving the chromatin boundary element SIS, and other 3′-RRs (Fig. 4a, upper and lower triangles).

**Fig. 4:**
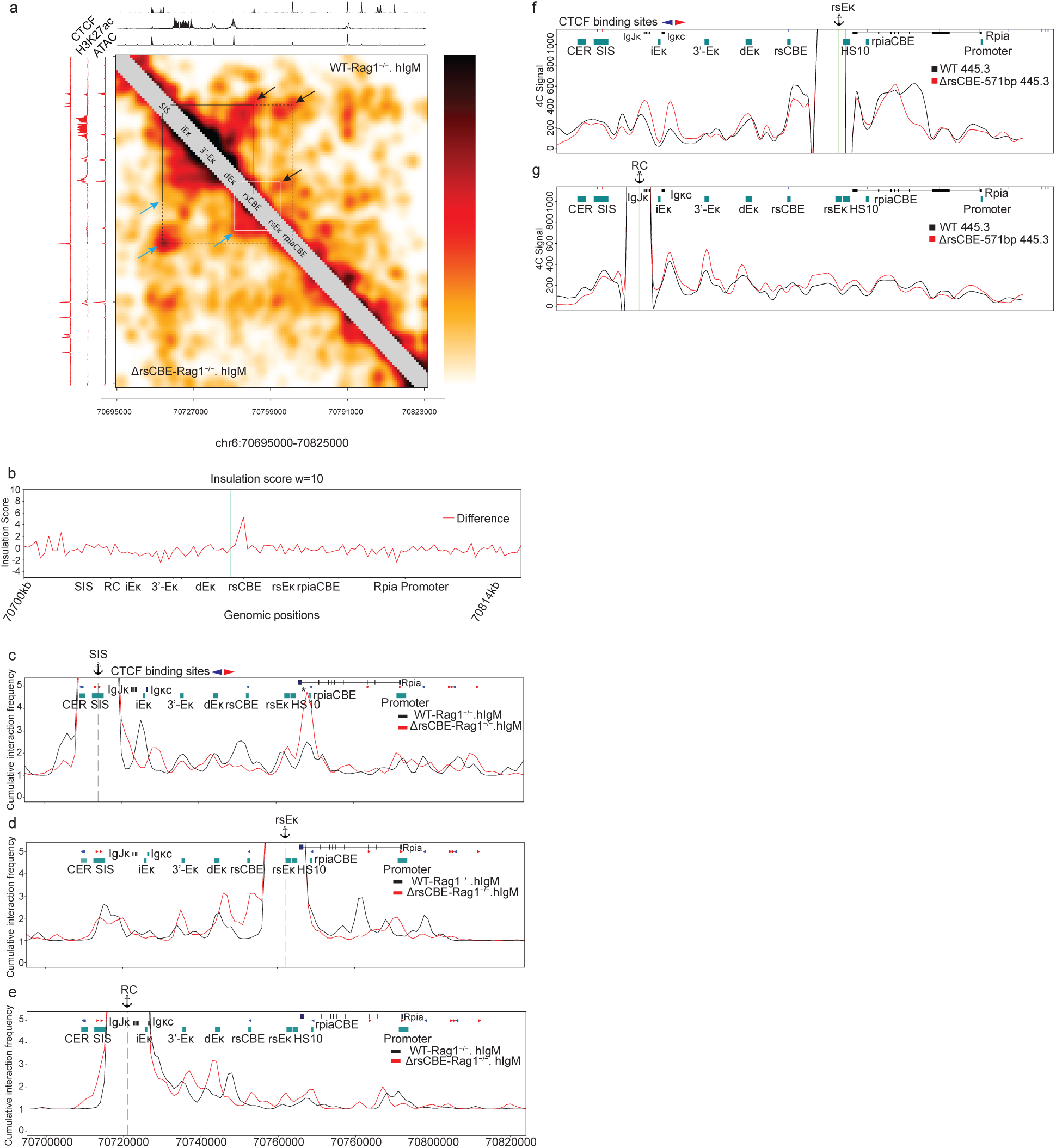
CBE controls the intricate Igκ super-enhancer architecture through chromatin insulation. **a,** High-resolution (1 kb) Micro-C-TALE contact maps of the 3′-RR, smoothed with a Gaussian filter (σ = 1 kb), overlaid with ATAC–seq, H3K27ac ChIP–seq, and CTCF ChIP– seq tracks. Notable interactions include SIS–rsCBE (black triangle), SIS–rpiaCBE (extended black triangle), and dEκ–rsEκ (white triangle). **b,** Insulation score profile plotted as a difference between ΔrsCBE-Rag1⁻/⁻.hIgM and WT-Rag1⁻/⁻.hIgM pre–B cells calculated with a 10-kb window. **c,** Virtual 4C profiles derived from Micro-C-TALE data (1-kb resolution; Gaussian σ = 1 kb) with SIS **d,** rsEκ, and **e,** RC as anchor viewpoints. **f,** 4C-seq interaction profiles across the 3′-RR with rsEκ, and **g,** RC as viewpoints in WT 445.3 and ΔrsCBE-571bp 445.3 cells. Interaction frequencies are shown as reads per million (RPM) with a 100-bp window and 21-bp running window. Data represent the mean of two biological replicates.

In the Micro-C-TALE interaction matrix, contact strength is indicated in the regions marked by black and blue arrows for WT-Rag1⁻/⁻.hIgM and ΔrsCBE-Rag1⁻/⁻.hIgM pre-B cells, respectively (Fig. 4a). Specifically, we compared chromatin interactions within the 3′-SEκ. In WT cells, contacts among the canonical enhancers were spatially constrained, primarily confined to the region between the SIS and rsCBE (Fig. 4a, upper black triangle), and, to a lesser extent, between the SIS and rpiaCBE (Fig. 4a, upper extended black triangle). The SIS–rsCBE domain encompasses the three canonical enhancers, thereby restricting their activity within this domain and limiting stronger interactions to the nearest downstream co-enhancers, such as HS10. By contrast, loss of rsCBE insulation in ΔrsCBE-Rag1⁻/⁻.hIgM pre-B cells led to pronounced changes in 3′-SEκ chromatin architecture. Contacts between the SIS and rsCBE were lost (Fig. 4a, lower black triangle), while spatial interactions within the SIS–rpiaCBE domain were enhanced (Fig. 4a, lower extended black triangle), permitting increased contacts in the region between the putative rsEκ shadow enhancer and dEκ (Fig. 4a, lower white triangle). These findings indicate that rsCBE functions as a key insulator, ensuring domain-specific chromatin interactions, thereby shaping enhancer connectivity and chromatin folding. Consistent with this altered chromatin topology, the rsEκ region exhibited elevated chromatin accessibility and H3K27ac enrichment in ΔrsCBE-Rag1⁻/⁻.hIgM pre-B cells (Fig. 3a,d).

The Igκ locus is organized into discrete sub-topologically associating domains (subTADs)^38,39,66^, demarcated by boundary elements enriched for architectural proteins such as CTCF and the cohesin subunit Rad21^30^. To assess the insulating capacity of the rsCBE region, we computed the Insulation Score (IS) in WT-Rag1⁻/⁻.hIgM and ΔrsCBE-Rag1⁻/⁻.hIgM pre-B cells. The IS of a locus measures the average interaction frequency between its upstream loci and downstream loci, such that dips in the IS reflect potential insulating regions^67^. Notably, deletion of rsCBE led to reduced insulation activity (higher IS) at this site, highlighting its essential role in preserving the 3′-SEκ chromatin architecture and subTAD boundary integrity (Fig. 4b).

To delineate the insulating function of rsCBE within the 3′-SEκ region further, we generated virtual 4C profiles from normalized Micro-C-TALE data at 1-kb resolution. Comparative analysis between WT-Rag1⁻/⁻.hIgM and ΔrsCBE-Rag1⁻/⁻.hIgM pre-B cells revealed that enhancer interactions across the 3′-SEκ were markedly altered upon rsCBE deletion. Using the SIS element as a virtual 4C viewpoint, we found that in WT-Rag1⁻/⁻.hIgM cells, the SIS interacts with rsCBE and rpiaCBE at comparable frequencies, consistent with published data (Fig. 4c)^40,68^. In ΔrsCBE-Rag1⁻/⁻.hIgM pre-B cells, loss of rsCBE led to aberrant chromatin looping toward the downstream rpiaCBE at higher frequency, incorporating the putative rsEκ shadow enhancer within the expanded interaction domain (Fig. 4c). Conversely, virtual 4C analysis from the rsCBE viewpoint revealed a loss of contact with the SIS element in ΔrsCBE-Rag1⁻/⁻.hIgM cells compared with WT-Rag1⁻/⁻.hIgM cells, indicating disruption of regular boundary activity (Extended Data Fig. 4b). Notably, the rsEκ viewpoint exhibited strengthened interactions with dEκ, 3′-Eκ, and the *Rpia* promoter in the absence of rsCBE (Fig. 4d). RC viewpoint analysis revealed enhanced contacts with canonical enhancers, including 3′-Eκ, dEκ, and the putative rsEκ shadow enhancer in ΔrsCBE-Rag1⁻/⁻.hIgM pre-B cells (Fig. 4e). Similarly, chromatin interactions from the rsCBE, 3′-Eκ, and dEκ viewpoints were evaluated (Extended Data Fig. 4b-d). These findings highlight a complex enhancer network driving elevated Igκ and *Rpia* expression in ΔrsCBE pre-B cells.

To reinforce our *in vivo* observations, we extended our analysis to the v-Abl– transformed pro-B cell line 445.3 to dissect the contribution of rsCBE to enhancer network organization within the 3′-SEκ. v-Abl–transformed lines are developmentally arrested at the pro-B to pre-B cell transition. Treatment of 445.3 WT cells with the Abl kinase inhibitor STI571 alleviates this block, inducing robust Igκ rearrangement and GLT^69^. The WT 445.3 line originates from a Rag1-deficient background, thereby preserving an intact germline Igκ locus and permitting the generation of deletion mutants without prior V(D)J recombination events. Using CRISPR–Cas9 genome editing, we deleted the 571-bp rsCBE region to recapitulate the *in vivo* deletion, hereafter referred to as ΔrsCBE-571bp 445.3, respectively. Single-cell clones were isolated and screened by DNA sequencing to confirm homozygous deletions. In particular, we investigated whether the putative rsEκ shadow enhancer functions as an active node within the reconfigured chromatin architecture that emerges following rsCBE deletion.

We conducted 4C-seq in the Rag1-deficient v-Abl pro–B cell line 445.3, enabling high-resolution mapping of chromatin contacts from a defined genomic viewpoint. Using rsEκ as a viewpoint in ΔrsCBE-571bp 445.3 cells, we observed increased contacts with IgκC and the RC compared with WT 445.3 cells, indicating that loss of rsCBE insulation promotes its integration into the canonical super-enhancer interaction hub. (Fig. 4f). To further explore this reorganization, we used RC as the reciprocal viewpoint in 4C-seq. In ΔrsCBE-571bp cells, the RC exhibited elevated interactions with 3′-Eκ, dEκ, rsEκ, and HS10 region, suggesting a substantial remodelling of the enhancer network upon loss of rsCBE insulation (Fig. 4g). When dEκ was used as the viewpoint, we noted reduced interactions with IgκC but maintained contact with the RC, alongside interaction with rsEκ (Extended Data Fig. 4e). These data indicate that rsCBE deletion leads to a reconfiguration of enhancer connectivity, with rsEκ emerging as a key regulatory node within the remodelled super-enhancer architecture. These findings reveal that rsCBE serves as a critical architectural boundary that orchestrates the structural integrity of the Igκ super-enhancer through chromatin insulation.

### rsCBE-mediated chromatin loop extrusion controls B-cell tolerance and autoimmunity

It is well established that murine λ⁺ B cells predominantly arise through receptor editing, which commences after secondary Igκ rearrangement^49,70–73^. The RS element, located approximately 25 kb downstream of the Cκ exon, harbours a canonical recombination signal sequence and undergoes V(D)J recombination with upstream Vκ segments or cryptic recombination sites within the Jκ–Cκ intron^49,74^. RS recombination results in the deletion of the Cκ exon and the three canonical Igκ enhancers, thereby functionally silencing the Igκ locus^49,71,72,75^. As RS rearrangement is predominant in murine λ⁺ B cells (77%)^76^ but occurs in only 12% of κ⁺ B cells^73^, we analysed Vκ–RS recombination in purified splenic λ⁺ B cells. Remarkably, Vκ–RS recombination was reduced approximately fivefold in ΔrsCBE mice compared to controls. Moreover, λ⁺ B cells from these mutants displayed decreased usage of Vκ– Jκ1 and enhanced usage of Vκ–Jκ5 segments, reflecting sequential κ locus rearrangements characteristic of receptor editing (Fig. 5a). This suggests that, in the absence of rsCBE, κ⁺ cells fail to transition into λ⁺ B cells, reflecting defective receptor editing^77^. In contrast, deletion of rpiaCBE alone had no detectable effect on Vκ–RS recombination (Fig. 5a). Collectively, these findings identify rsCBE as a key regulator of Vκ–RS recombination and an essential determinant of λ⁺ B cell development.

**Fig. 5:**
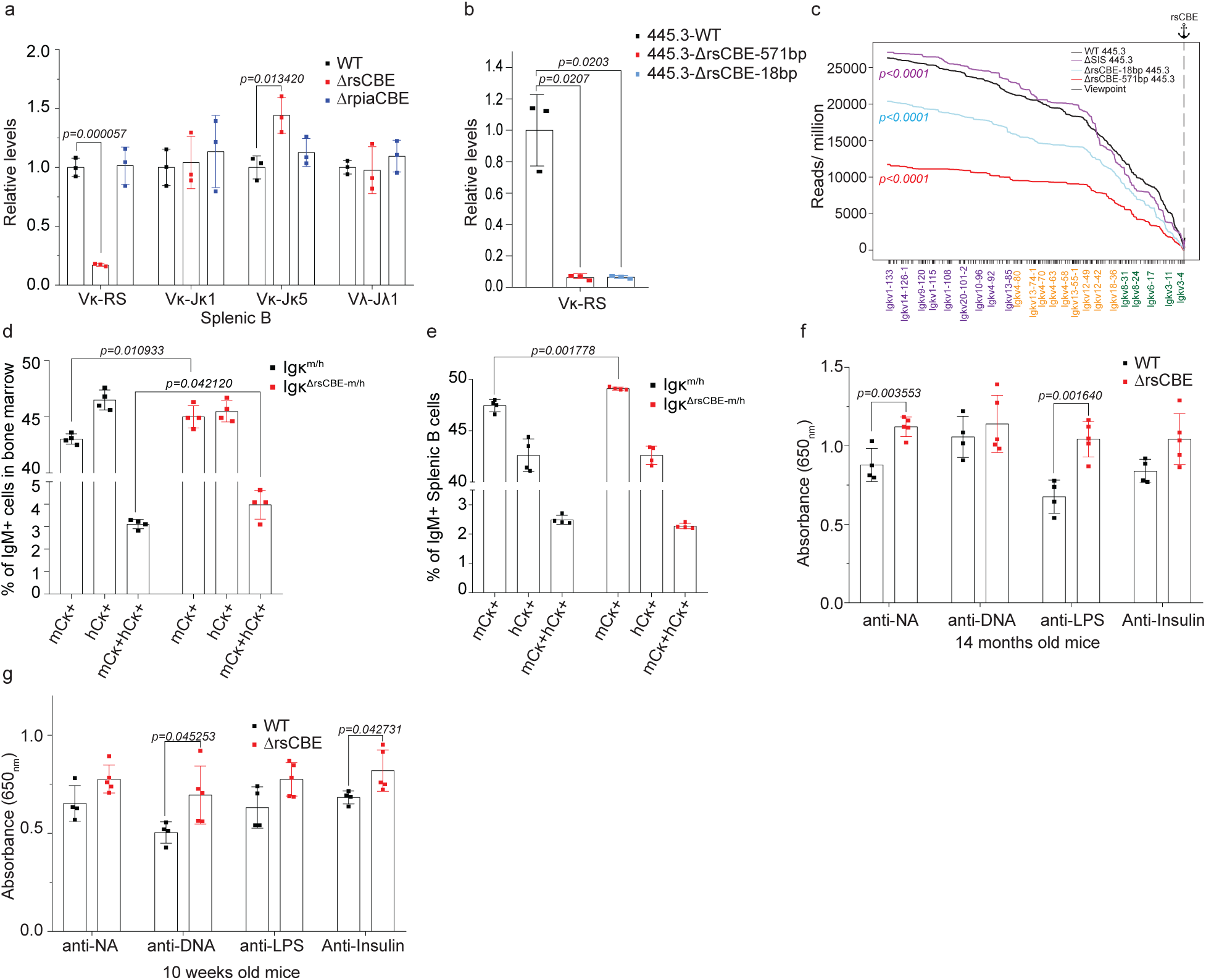
rsCBE-mediated chromatin loop extrusion control B-cell tolerance and autoimmunity. **a,** Relative levels of rearranged Jκ, Jλ1, and Vκ–RS segments in FACS-purified B220⁺Igλ⁺ splenic B cells, quantified by qPCR and normalized to the Eμ genomic region (n = 3; mean ± s.d.). **b,** Relative Vκ–RS recombination in Rag1-proficient WT 445.3, ΔrsCBE-571 bp 445.3, and ΔrsCBE-18 bp 445.3 clones, measured by qPCR and normalized to the Eμ genomic region (n = 3). P values were determined by unpaired t-test. **c,** Cumulative frequency distributions (CFDs) from rsCBE viewpoints showing cumulative 4C signals in ΔrsCBE-571 bp, ΔrsCBE-18 bp, and ΔSIS 445.3 cells relative to WT 445.3 (p < 0.0001, Mann– Whitney test). Data represent the mean of two independent biological replicates. **d,** Allelic exclusion in bone marrow IgM⁺ B cells from Igκ^m/h^ and Igκ^ΔrsCBE-m/h^ mice, shown as percentages of mCκ⁺, hCκ⁺, and dual mCκ⁺hCκ⁺ cells (n = 4). **e,** Allelic exclusion in splenic IgM⁺ B cells from Igκ^m/h^ and Igκ^ΔrsCBE-m/h^ mice, shown as percentages (n = 4). Data are mean ± s.d.; P values were determined by unpaired t-test. **f,** Serum IgG autoantibodies against nuclear antigens (NA), DNA, LPS, and insulin quantified by ELISA in 14-month-old mice (n = 4–10) and **g,** 10-week-old mice (n = 4–5), measured by ELISA. Data are mean ± s.d.; P values were determined by unpaired t-test.

To investigate whether CTCF-mediated chromatin loop extrusion drives Vκ–RS recombination, we focused on the v-Abl–transformed pro-B cell line. We generated an additional deletion targeting the 18 bp CTCF-binding motif within rsCBE, hereafter referred to as ΔrsCBE-18bp 445.3, respectively. Single-cell clones were isolated and screened by DNA sequencing to confirm homozygous deletions. Additionally, the ΔrsCBE-18bp clones were validated by ChIP-qPCR to confirm loss of CTCF binding (Extended Data Fig. 5a). Given that 445.3 cells are Rag1-deficient, both the 445.3 WT and rsCBE-deleted lines were transduced with a Rag1-expressing retrovirus to restore recombination capability. Following transduction, cells were treated with STI571 for 20 hours, and Vκ–RS recombination levels were quantified by qPCR. Remarkably, the ΔrsCBE-18bp 445.3 clone lacking the CTCF-binding motif recapitulated the reduction in Vκ–RS recombination observed with the larger ΔrsCBE-571bp deletion, both *in vitro* and *in vivo*. (Fig. 5b). These results strongly support the model that CTCF-mediated chromatin loop extrusion at the rsCBE locus is a critical driver of Vκ–RS recombination, facilitating the development of λ⁺ B cells.

Primary rearrangements, whether by deletion or inversion, typically delete or invert the CER and SIS elements^40^. *In vitro* studies with v-Abl pro-B cells suggest that RAG scanning from the primary Jκ–RC is halted by the CTCF-dependent SIS element^40^. We therefore propose that the SIS element acts as a barrier to rsCBE-mediated CTCF interactions with the Vκ region. Consequently, deletion and inversion of CER and SIS during primary rearrangement may enable rsCBE to interact with the Vκ region, thereby promoting Vκ–RS secondary recombination.

To investigate long-range chromatin interactions between rsCBE and Vκ gene segments, and to assess whether the SIS functions as a barrier to rsCBE-mediated CTCF contacts with the Vκ region, we performed *in vitro* studies using the v-Abl– transformed pro-B cell line. Using 4C-seq with a viewpoint positioned upstream of rsCBE, we found that rsCBE establishes contacts with dEκ, 3′-Eκ, IgκC, RC, SIS, and the CER element in WT 445.3 cells (Extended Data Fig. 5b). By contrast, both ΔrsCBE-571bp and ΔrsCBE-18bp 445.3 cells showed markedly reduced interactions with these regulatory elements, with the most pronounced loss observed for contacts with SIS (as expected) and CER, (Extended Data Fig. 5b). To quantitatively assess alterations in long-range chromatin interaction, we performed cumulative frequency distribution (CFD) analysis of 4C-seq read density upstream of the rsCBE viewpoint. This approach allowed us to systematically measure the extent of rsCBE–Vκ region interactions across the locus. Strikingly, both the frequency and intensity of rsCBE– Vκ contacts were markedly diminished in ΔrsCBE-571bp and ΔrsCBE-18bp 445.3 cells, with the ΔrsCBE-571bp deletion exhibiting an even more pronounced defect, underscoring the critical role of rsCBE in maintaining productive long-range interactions with distal Vκ gene segments. (Fig. 5c).

To explore chromatin interactions during secondary rearrangement, we also deleted the ∼3 kb SIS element, which harbours two CTCF-binding sites oriented toward rsCBE, and is thought to constrain rsCBE–Vκ interactions in the v-Abl–transformed line. Homozygous ΔSIS 445.3 single-cell clones were confirmed by DNA sequencing and used for downstream assays. As shown above, 4C-seq analysis revealed that in WT 445.3 cells, rsCBE engages in interactions with dEκ, 3′-Eκ, IgκC, RC, SIS, and the CER element (Extended Data Fig. 5c). By contrast, in ΔSIS 445.3 cells, rsCBE exhibited a pronounced loss of contact with IgκC and SIS, accompanied by enhanced interactions with the Vκ region, consistent with SIS functioning as a chromatin barrier that modulates rsCBE–Vκ contact dynamics (Extended Data Fig. 5c). In ΔSIS 445.3 cells, CFD analysis revealed a marked increase in upstream interactions, toward the Vκ region, compared to WT 445.3 cells (Fig. 5c). Indeed, deletion of the SIS element *in vivo* has been shown to increase Vκ–RS recombination by ∼1.2-fold in splenic λ⁺ B cells^22^. These findings suggest that SIS constrains rsCBE activity during primary rearrangement. At the same time, rsCBE promotes chromatin looping toward the Vκ gene cluster to drive secondary rearrangements, when the SIS element is absent.

We subsequently investigated how deletion of rsCBE influences allelic exclusion, with particular emphasis on assessing the emergence of dual κ-expressing B cells in ΔrsCBE mice. Given that allelic inclusion is a consequence of receptor editing, it may facilitate the emergence and persistence of autoreactive B cells by circumventing central tolerance mechanisms, thereby contributing to autoimmunity^78–82^. To evaluate this, we crossed ΔrsCBE with human Cκ knock-in mice to generate Igκ^m/h^ heterozygotes and analysed IgM⁺ immature B cells in the bone marrow (Extended Data Fig. 5d). Flow cytometric analysis revealed that rsCBE deficiency resulted in a modest yet significant increase in the proportion of cells expressing the mouse Cκ (mCκ) allele, accompanied by a modest, non-significant decrease in human Cκ (hCκ). (Fig. 5d, Extended Data Fig. 5e). Notably, ΔrsCBE heterozygotes also exhibited a significant increase in the frequency of dual-expressing mCκ⁺hCκ⁺ cells, indicative of allelic inclusion (Fig. 5d). These findings suggest that rsCBE promotes preferential usage of the Igκ allele in cis and plays a role in enforcing allelic exclusion during early B cell development.

We further assessed allelic usage in the spleen of Igκ^m/h^ heterozygous mice. Flow cytometric analysis of splenic B220⁺ B cells confirmed that loss of rsCBE resulted in a modest but significant increase in the proportion of cells expressing the mCκ allele (Fig. 5e, Extended Data Fig. 5e). Our data demonstrate that deletion of rsCBE results in a small breakdown of allelic exclusion in the bone marrow, as evidenced by the emergence of dual Igκ allele-expressing (mCκ⁺hCκ⁺) immature B cells.

Receptor editing is a key mechanism of B cell tolerance, whereby κ⁺ B cells first attempt secondary κ rearrangements to replace an autoreactive κ chain with a non-autoreactive one. If autoreactivity persists, RS recombination is triggered to activate the λ locus and express a λ light chain, thereby restoring self-tolerance^49^. Inefficient RS rearrangement can compromise this process, allowing autoreactive B cells to persist in the periphery, a feature observed in murine models of systemic lupus erythematosus (SLE) and type 1 diabetes (T1D), as well as in patients with SLE^83,84^. Given that RS rearrangement is diminished in such models, we hypothesized that impaired RS recombination in splenic λ⁺ B cells from ΔrsCBE mice may underlie defects in central B cell tolerance. Thus, we measured serum levels of autoantibodies targeting canonical self-antigens^85–88^ in ΔrsCBE mice. Notably, ΔrsCBE mice exhibited elevated titres of IgG-class anti-nuclear antigen (ANA) and anti-LPS antibodies at 14 months of age, as assessed by ELISA—hallmarks of systemic autoimmune pathology (Fig. 5f). Serum levels of anti-DNA and anti-insulin also showed an elevated trend compared to wild-type controls (Fig. 5f). Analysis of splenic B cell subsets in ΔrsCBE mice revealed no major alterations, although the mice exhibited splenomegaly, indicating inflammation (Extended Data Fig. 5f,g). These findings collectively suggest that loss of rsCBE compromises immune tolerance and precipitates autoantibody production.

To examine whether elevated Igκ expression in ΔrsCBE mice contributes to early-onset of systemic-like autoimmunity, we assessed serum autoantibody levels at 10 weeks of age. Notably, ΔrsCBE mice displayed an increase in IgG autoantibodies targeting DNA and insulin compared to wild-type controls (Fig. 5g). These findings implicate dysregulated Igκ expression, driven by the loss of rsCBE insulation, in possibly promoting early autoimmune responses. These findings underscore the regulatory importance of enhancer insulation in preserving B-cell immune tolerance from a young age.

## Discussion

The RS element is a central regulator of Igλ expression, light-chain isotype exclusion, and B cell receptor editing^49^, yet the epigenetic basis of these processes has remained elusive. Here, we show that rsCBE-mediated chromatin loop extrusion establishes a CTCF-dependent axis that mechanistically underpins this regulation. Loss of rsCBE insulation unmasks a previously sequestered putative shadow enhancer, rsEκ, which aberrantly integrates into the 3′-SEκ regulatory hub to drive coordinated misregulation of Igκ and the closely linked gene *Rpia*. This architectural disruption not only enhances Igκ expression but also predisposes to autoantibody production, directly linking chromatin topology to the maintenance of central B cell tolerance. More broadly, our findings demonstrate that individual cis-regulatory CBEs within the 3′-SEκ function as distinct, bidirectional transcriptional insulators that safeguard the integrity of transcription.

Our findings establish insulation within the 3′-SEκ as a critical determinant of gene regulation in B cells. Loss of rsCBE produced a ∼1.8-fold increase in Igκ expression in pre-B cells, whereas rpiaCBE deletion caused a modest but significant reduction. Strikingly, both deletions derepressed the neighbouring *Rpia* gene, demonstrating that individual CBEs within the 3′-SEκ function as non-redundant architectural boundaries that differentially tune enhancer–promoter communication. This mechanistic insight contrasts with earlier reports showing that deletion of SIS, CER, or both elements within the 3′-SEκ had minimal impact on Igκ or *Rpia* expression, leaving unresolved how *Rpia* was insulated from the potent regulatory activity of the 3′-SEκ^11,22,42–44^. Even loss of HS10, the co-enhancer closest to *Rpia*, failed to perturb its expression^14^, underscoring that enhancer proximity alone does not dictate insulation. Furthermore, CTCF gene deletion in pre-B cells was shown to reduce both 5′- and 3′-GLTs at the Jκ region while increasing proximal Vκ GLT and elevating GLTs at the Jλ1 and Jλ3 regions^20^. Similarly, mutations in Igκ enhancers within the 3′-SEκ decrease Igκ expression but enhance Igλ recombination^13,18,74,89,90^. The simultaneous upregulation of Igκ and *Rpia*, or of either gene individually, has not been observed in previous studies, emphasizing the unique role of rsCBE-mediated insulation in preserving balanced gene regulation within B cells. Given that aberrant *Rpia* activation has been implicated in autophagy and diverse cancers^53–56^, our findings highlight CBE-dependent chromatin architecture as an essential mechanism that safeguards *Rpia* insulation within the 3′-SEκ. Whether the increased expression of the *Rpia* gene affects B cell function or survival warrants further investigation.

Our study identifies rsCBE as a critical architectural element in early B cell development, functioning to insulate the putative rsEκ shadow enhancer and prevent its premature activation. Loss of rsCBE insulation in adult mice most probably unleashes rsEκ activity in pro-B cells, driving untimely Igκ recombination. This premature recombination imposes a G1–S checkpoint arrest during the pro-B to immature B cell transition, resulting in attrition of the pre-B cell pool through apoptosis. In earlier studies, Igκ rearrangement was induced in fetal-derived B1 pro-B cells^91^. Thus, rsCBE safeguards developmental timing by constraining enhancer activity until the appropriate stage, and its loss reveals that Igκ recombination can be induced in adult pro-B cells.

Loss of rsCBE and rpiaCBE insulation triggers profound epigenetic remodeling within the 3′-SEκ. Disruption of rsCBE enhances chromatin accessibility at rsEκ, HS10, and rpiaCBE, while concomitantly reducing accessibility at the RS element in pre-B cells. This suggests that rsCBE functions as a gatekeeper of Rag1 accessibility, with CTCF-mediated loop extrusion directing Vκ–RS synapsis and recombination^21^. The accompanying increase in H3K27ac across rsEκ, HS10, 3′-Eκ, and the RC, together with enrichment at Vκ RSSs and Vκ genes, indicates that enhancer activation and chromatin remodelling are tightly coupled to recombination potential^63–65^. Thus, rsCBE insulation safeguards the balance between enhancer-driven transcriptional activation and recombinase accessibility, and its loss disrupts this architecture to prematurely couple Igκ expression with aberrant Vκ–RS recombination in pre-B cells.

The 3′-SEκ functions as a highly organized regulatory hub of enhancer interactions, integrating the CER, SIS, rsCBE, and rpiaCBE into its chromatin architecture^37^. Within the 3′-SEκ network, rsCBE and SIS CBE form a boundary domain that encapsulates the three canonical enhancers, thereby constraining their activity to maintain physiological levels of Igκ and *Rpia* expression. Disruption of rsCBE insulation rewires this architecture by looping toward rpiaCBE and incorporating the previously sequestered putative rsEκ shadow enhancer, leading to simultaneous upregulation of both Igκ and *Rpia*. Beyond transcriptional control, rsCBE is indispensable for Vκ–RS recombination in pre-B cells, akin to the RS element^49^. Loss of rsCBE reduces RS element accessibility *in vivo* and abrogates Rag1-mediated recombination. This is in line with an established study showing that CBEs mediate accessibility of Rag substrates during chromatin scanning^21^. It is also consistent with models of RAG scanning, which propose that RAG complexes search for RSSs located within convergent CTCF sites in the IgH locus^92–99^.

Functionally, increased Igκ expression skews light-chain isotype usage, producing a twofold reduction in λ⁺ cells, due to impaired receptor editing and hyperactivation of the Igκ locus. Interestingly, *in vitro* analyses demonstrate that rsCBE facilitates Vκ– RS secondary rearrangements through direct interactions with the Vκ variable region, contacts that are normally constrained by the SIS element^40^. Importantly, primary rearrangements involving deletion or inversion of CER and SIS remove this barrier, allowing rsCBE to engage during secondary editing. Together, these findings establish rsCBE as a dual-function regulator that coordinates enhancer insulation, Vκ–RS recombination, and light-chain isotype balance.

While light chain isotype exclusion remained intact, allelic exclusion was compromised. Hybrid reporter mice revealed an increased frequency of immature B cells co-expressing both Igκ alleles (mCκ⁺hCκ⁺), demonstrating a requirement for rsCBE in maintaining allelic exclusion. The presence of such dual-mCκ⁺hCκ⁺ cells in the bone marrow suggests defective clonal deletion, possibly because of dilution of autoreactive BCRs^78–80^. An additional layer of peripheral clonal deletion is still active and effective, as dual-κ+ cells are absent in the periphery. However, this peripheral deletion mechanism is not absolute, as evidenced by increased autoantibody titres in the serum. These findings establish rsCBE as a cis-regulatory element that safeguards B cell tolerance. By coordinating receptor editing and allelic exclusion, rsCBE prevents the emergence of autoreactive clones and preserves immune homeostasis. Its loss uncovers a critical checkpoint in central tolerance, where Igκ hyperactivation and defective editing converge to accelerate the generation of autoreactive B cells.

B cells are subject to stringent selection in the bone marrow to eliminate autoreactive clones as part of the central tolerance mechanism. Such cells can undergo receptor editing, clonal deletion, or anergy to generate a non-autoreactive repertoire. Deletion of rsCBE *in vivo* impaired receptor editing, leading to a reduction in λ⁺ B cells and spontaneous production of IgG autoantibodies as early as ten weeks of age. Notably, rsCBE-deficient mice exhibited serum autoantibodies against DNA and insulin— hallmarks of systemic autoimmune disease such as lupus, well before they typically arise in C57BL/6 mice, which normally do not develop anti-DNA antibodies until late in life^100^. These findings indicate that rsCBE is required to preserve B-cell tolerance at a young age. At 14 months of age, rsCBE-deficient mice displayed elevated IgG autoantibody titres against nuclear antigens (ANA) and lipopolysaccharide (LPS), further underscoring rsCBE long-term role in immune homeostasis.

Previous studies have shown that germline RS mutations reduce the frequency of λ⁺ B cells by impairing receptor editing, while leaving Igκ rearrangement and allelic exclusion intact^49^. In contrast, rsCBE deletion not only recapitulates editing defects but also drives excessive Igκ rearrangement in pre-B cells. This shift skews light chain usage, with splenic λ⁺ B cells showing increased detection of Vκ–Jκ5 segments, reflecting a failure to undergo Vκ–RS recombination in the absence of rsCBE, a defining feature of secondary rearrangements. The marked reduction in λ⁺ cells, coupled with increased κ⁺ cells in bone marrow and spleen, highlights the essential role of rsCBE in sustaining λ⁺ B cell development.

Transgenic mouse models of defective receptor editing manifest enhanced clonal deletion without spontaneous autoantibody production^101–103^. Even under conditions that block apoptosis, such as Bcl-2 overexpression, inefficiently edited B cells generally remain subject to tolerance mechanisms with minimal autoantibody production^104–106^. Only when the RS mutation is combined with anti-apoptotic signals has autoimmunity been observed, and then only in non-physiological settings^49,105–107^. By contrast, loss of rsCBE alone is sufficient to disrupt receptor editing and trigger early breakage of B cell tolerance. Unlike transgenic models of central tolerance, which express a high-avidity monoclonal BCR that primarily promotes clonal deletion and peripheral anergy, rsCBE-deficient mice retain a normal polyclonal BCR repertoire. This physiological diversity reveals a broader escape from both central and peripheral tolerance checkpoints.

Our findings position rsCBE as a dual-function regulatory element that couples chromatin architecture with B cell tolerance. By insulating the putative rsEκ shadow enhancer and directing CTCF–cohesin loop extrusion, rsCBE safeguards the developmental timing of Igκ recombination, maintains allelic exclusion, and preserves the balance of light-chain isotypes. Its loss disrupts enhancer insulation, deregulates Igκ and *Rpia* expression, and impairs Vκ–RS recombination, thereby compromising receptor editing. The early production of autoantibodies in rsCBE-deficient mice at a young age directly links chromatin architecture to central tolerance. Our study demonstrates that defects in central B cell tolerance predispose to failures in peripheral tolerance, culminating in early autoantibody production. More broadly, our study demonstrates that individual CBEs within the 3′-SEκ act as non-redundant architectural boundaries that orchestrate enhancer cross-talk and promoter engagement to preserve transcriptional integrity.

## Methods

### Mice

Transgenic mice expressing a human immunoglobulin heavy chain (Igh-Tg⁺), originally described, were generously provided by Dr. Cornelis Murre (University of California, San Diego)^108^. Mice carrying a human Cκ knock-in gene were kindly provided by Michel C. Nussenzweig (The Rockefeller University, New York, NY)^78^. All animal procedures were approved by the Institutional Animal Care and Use Committee of the Hebrew University of Jerusalem (MD-24-17515-4). Wild-type C57BL/6 and genetically modified mice were housed under specific pathogen-free (SPF) conditions. Animals were maintained on a 12-hour light/dark cycle at 22 ± 2 °C and 55 ± 15% relative humidity. Unless otherwise indicated in the figure legends, male mice aged 6–12 weeks were used for experiments. All lines were maintained on a C57BL/6 genetic background. For the allelic exclusion experiments, reporter mice were maintained on a B6/BALB/c hybrid background.

### Generation of rsCBE and rpiaCBE deletions in mice using CRISPR–Cas9

To generate an 18-bp rsCBE deletion in 445.3 cells, a single-stranded oligodeoxynucleotide (ssODN; IDT) containing 50-bp homology arms flanking the 18-bp CTCF-binding motif was used as a homology-directed repair (HDR) template. A single-guide RNA (sgRNA) targeting the CTCF-binding motif within the 571-bp rsCBE region (rsCBE-sgRNA3) was designed using the CRISPR design tool (http://crispr.mit.edu) and cloned into pSpCas9(BB)-2A-GFP (pX458; Addgene #48138). To generate the 571-bp rsCBE deletion, two sgRNAs (rsCBE-sgRNA1 and rsCBE-sgRNA2) were individually cloned into pX458, and all constructs were sequence-verified. The ∼3 kb SIS deletion was produced using two sgRNAs (SIS-sgRNA1 and SIS-sgRNA2), which were likewise cloned and verified. pX458 carrying rsCBE-sgRNA3 together with the ssODN were co-transfected into 445.3 cells using the Neon electroporation system to induce the 18-bp deletion, whereas the 571-bp deletion was achieved by co-transfecting the two rsCBE pX458 constructs. Similarly, the SIS pX458 pair was co-transfected to generate the ∼3 kb deletion. After 48 h, GFP⁺ cells were single-cell sorted by FACS and cultured for two weeks to allow colony formation. Individual clones were screened by PCR for biallelic deletions, and successful editing was confirmed by Sanger sequencing. CRISPR/Cas9-mediated genomic modifications were validated by PCR and DNA sequencing.

### Cells

Bone marrow-derived cells were cultured on an ST2 stromal feeder layer (kindly provided by A. Rolink)^109^ following density gradient separation using Histopaque. Cells were maintained in RPMI 1640 medium supplemented with 10% fetal bovine serum (FBS), 2 mM L-glutamine, 100 U/ml penicillin, 100 μg/ml streptomycin, and 50 μM 2-mercaptoethanol. Interleukin-7 (IL-7; PeproTech, Cat. 217-17) was added to the culture at a final concentration of 5 ng/ml. The Abelson virus-transformed pro-B cell line 445.3 was maintained in RPMI 1640 medium supplemented with 10% FBS, 2 mM L-glutamine, 100 U/ml penicillin, 100 μg/ml streptomycin, and 50 μM 2-mercaptoethanol. The 445.3 cells are Rag1−/− and were kindly provided by Dr. Cornelis Murre (UCSD). Hep-2 cells were also cultured in RPMI 1640 medium supplemented with 10% FBS, 2 mM L-glutamine, 100 U/ml penicillin, 100 μg/ml streptomycin, and 50 μM 2-mercaptoethanol. HEK 293T cells were maintained in DMEM supplemented with 10% FBS, 2 mM L-glutamine, and 100 U/ml penicillin–100 μg/ml streptomycin.

### Bone marrow and splenic B cell extraction

The femur, tibia, and fibula (and humeri in some instances) were dissected, rinsed with PBS, and mechanically homogenized in 2 ml of FACS buffer (0.5% BSA in PBS) using a mortar and pestle until completely depigmented. The resulting cell suspension was filtered through a 100 μm nylon mesh into a 50 ml conical tube and centrifuged at 300 × g for 5 min at 4 °C. The cell pellet was subsequently resuspended in an appropriate volume of FACS buffer for downstream use. Lymphocytes were enriched by layering cell suspensions onto Histopaque-1077 (Sigma-Aldrich), followed by centrifugation at 700 × g for 30 min at 4 °C without brake. The interface containing enriched lymphocytes was carefully collected, washed with FACS buffer, and pelleted by centrifugation at 1,200 rpm for 5 min at 4 °C. The cell pellet was resuspended in 300 µl of chilled FACS buffer and labelled for flow cytometric sorting. For flow cytometric analysis, bone marrow and splenic cells were treated with 1× Ammonium–Chloride–Potassium (ACK) lysis buffer to remove red blood cells, then washed and resuspended in an appropriate volume for analysis.

### Flow Cytometry

Pre-B cells (B220^+^IgM^-^CD43^-^CD25^+^) were isolated from the bone marrow of pooled cohorts of five to six female mice (6–12 weeks old) for Igκ RNA library preparation. DNA and RNA were extracted from samples of individual mice for subsequent qPCR analysis. Bone marrow cell suspensions were counted, washed in 10 ml of phosphate-buffered saline (PBS), and resuspended at 1 × 10⁶ cells per 100 μl in FACS buffer containing Fc receptor blocking reagent (anti-CD16/CD32, Invitrogen, #14-0161-82; 1:100 dilution). Cells were stained for 25 min at 4 °C with the following antibodies (1:200–1:400): anti-B220 (RA3-6B2, PerCP/Cy5.5, BioLegend, #103235); anti-IgM (II/41, APC, eBioscience, #17-5790-82; II/41, eFluor™ 450, Invitrogen, #48-5790-82); anti-CD25 (PC61.5, Alexa Fluor 488, Invitrogen, #53-0251-82; PC61, BV786, BD Horizon, #564023); anti-CD19 (1D3/CD19, PE-Cy7, BioLegend, #152417; 1D3/CD19, APC, BioLegend, #152410); and anti-CD43 (eBioR2/60, PE, Invitrogen, #12-0431-81). After staining, cells were washed and resuspended in FACS buffer for analysis on a Beckman CytoFLEX cytometer. Data were analysed using FlowJo v10.7 (BD Biosciences), and cells were sorted on a BD FACSAria.

For splenic B cell analysis, single-cell suspensions were first incubated with Fc receptor–blocking antibody for 10 min at 4 °C. After washing, cells were stained for 25 min at 4 °C with the following antibodies: anti-B220; anti-IgM (II/41, eFluor™ 450, Invitrogen, #48-5790-82); anti-CD93 (AA4.1, APC, eBioscience/Invitrogen, #17-5892-81); anti-CD21 (7G6, FITC, BD Pharmingen, #561769); and anti-CD23 (B3B4, PE, BD Pharmingen, #561773).

For analysis of immunoglobulin light chain isotypes, splenic single-cell suspensions were stained for 25 min at 4 °C with anti-mouse Cκ (H139-52.1, PE, SouthernBiotech, #1180-09), anti-Igλ1–3 (R26-46, BD Pharmingen, #553434), and anti-B220. For bone marrow immature B cell analysis, cells were stained with anti-CD19 (1D3/CD19, PE-Cy7, BioLegend, #152417), anti-B220, anti-IgM (II/41, APC, eBioscience, #17-5790-82), as well as anti-mouse Cκ and anti-Igλ1–3.

For assessment of allelic exclusion, single-cell suspensions were stained with anti-IgM (II/41, eFluor™ 450, Thermo Fisher #42-5790-82) in combination with anti-mouse Cκ. After washing, cells were further incubated with anti-human Cκ (SB81a, APC, SouthernBiotech, #9230-11), followed by a final wash and filtration through a 40 μm cell strainer. Samples were acquired on a flow cytometer and analysed using FlowJo v10.7 (BD Biosciences).

### CRISPR-Cas9 mediated mutagenesis

To generate an 18-bp rsCBE deletion in 445.3 cells, a single-stranded oligodeoxynucleotide (ssODN; IDT) containing 50-bp homology arms flanking the 18-bp CTCF-binding motif was used as a homology-directed repair (HDR) template. A single-guide RNA (sgRNA) targeting the CTCF-binding motif within the 571-bp rsCBE region (rsCBE-sgRNA3) was designed using the CRISPR design tool (http://crispr.mit.edu) and cloned into pSpCas9(BB)-2A-GFP (PX458; Addgene #48138). To generate the 571-bp rsCBE deletion, two sgRNAs (rsCBE-sgRNA1 and rsCBE-sgRNA2) were individually cloned into PX458, and all constructs were sequence-verified. The ∼3 kb SIS deletion was produced using two sgRNAs (SIS-sgRNA1 and SIS-sgRNA2), which were likewise cloned and verified. PX458 carrying rsCBE-sgRNA3 together with the ssODN were co-transfected into 445.3 cells using the Neon electroporation system to induce the 18-bp deletion, whereas the 571-bp deletion was achieved by co-transfecting the two rsCBE PX458 constructs. Similarly, the SIS PX458 pair was co-transfected to generate the ∼3 kb deletion. After 48 h, GFP⁺ cells were single-cell sorted by FACS and cultured for two weeks to allow colony formation. Individual clones were screened by PCR for biallelic deletions, and successful editing was confirmed by Sanger sequencing. CRISPR/Cas9-mediated genomic modifications were validated by PCR and DNA sequencing.

### RNA and genomic DNA analysis

Total RNA was extracted from 4–6 × 10⁵ FACS-purified cells using the High Pure RNA Isolation Kit (Roche, #11828665001) or the Nucleospin RNA Kit (Macherey-Nagel, #740955.50) according to the manufacturers’ instructions. First-strand cDNA was synthesized from 30–60 ng of total RNA using the qScript cDNA Synthesis Kit (Quanta Bio, #95047-100). Genomic DNA was isolated from a similar number of cells using the DNeasy Blood & Tissue Kit (Qiagen, #96504). Quantitative real-time PCR (qPCR) was performed on a Bio-Rad CFX Connect system using SYBR Green Master Mix.

### 4C Sequencing

4C template and library preparation was performed as described previously^110^. In brief, 5–10 × 10⁶ Rag1⁻/⁻ 445.3-WT, ΔrsCBE-571 bp 445.3, ΔrsCBE-18 bp 445.3, and ΔSIS 445.3 cells were crosslinked with 1% formaldehyde following stimulation with kinase inhibitors. Primary digestion was performed using NlaIII (NEB, R0125S), followed by dilution and ligation. Cross-links were then reversed by proteinase K treatment, and templates were trimmed with Csp6I (NEB, R0639S) before re-ligation. Inverted PCR was carried out from multiple viewpoints using the Expand Long Template PCR System (Roche, #11681842001) with indexed Illumina sequencing adapters, and products were purified using the Qiaquick PCR Purification Kit (Qiagen, #28104). Libraries were sequenced on an Illumina NovaSeq 6000 with 122–160 bp single-end reads. 4C-Seq data were analysed using the pipe4C pipeline^110^, and genomic interaction profiles were visualized by smoothing WIG files with the npreg R package (spar = 0.75). 4C data were normalized to reads per million (RPM). CFD differences upstream of the rsCBE viewpoint among WT 445.3, ΔrsCBE-571 bp 445.3, ΔrsCBE-18 bp 445.3, and ΔSIS 445.3 were assessed for significance using the Mann–Whitney test in Prism 8.0.1.

### Retrovirus preparation and transduction

To generate a Rag1-expressing retrovirus, HEK 293T cells at 70–80% confluence were co-transfected with pMSCV-IRES-Bsr-Rag1, encoding full-length wild-type murine Rag1 and a blasticidin resistance cassette, and the pEco packaging plasmid, which expresses the ecotropic envelope protein under the CMV immediate-early promoter. Transfections were performed using Mirus reagent according to the manufacturer’s instructions, and viral supernatants were collected 48 h post-transfection. For *in vitro* recombination assays, 445.3 pro-B cells were transduced with the Rag1 retrovirus and cultured in blasticidin (20 µg/mL) for seven days to select for stable expression. Cells were subsequently treated with STI571 to induce recombination.

### Igκ repertoire RNA sequencing and analysis

DNase-treated RNA was isolated from FACS-purified pre-B cells from bone marrow. The library preparation protocol was adapted from Rena Levin-Klein et al. 2017^111^. RNA was poly-A enriched using poly-dT beads (Life Technologies) in two selection cycles and RT then performed using AffinityScript QPCR cDNA Synthesis Kit (Agilent) with an RT primer specific for the Cκ region 5′-ATGCTGTAGGTGCTGTCTTT-3′. The residual RNA was degraded with 0.1 N NaOH, neutralized with 0.1 M acetic acid and the single strand cDNA then purified using Silane beads (Life Technologies). A 3TR3 adapter (5′-/Phos/ AGATCGGAAGAGCACACGTCTG/3SpC3/-3′) was ligated to the 3′ end by overnight incubation with T4 RNA ligase (NEB) at 22 °C, and the cDNA then purified from excess adapter with Silane beads and PCR amplified for 12 cycles using the reverse complement of the 3RT3 adapter as the forward primer and the upstream Cκ region as the reverse primer with the partial Truseq Illumina adapter added to the beginning (5′-TACACGACGCTCTTCCGATCT-ACTGGATGGTGGGAAGATGGAT-3′). The PCR product was cleaned with 0.7 × ampure XT beads, amplified with indexed universal Illumina adapter primers for an additional seven cycles to obtain ∼550 bp libraries. These libraries were sequenced on a Miseq platform yielding 151 bp paired-end. Sequencing reads were processed to retain only read pairs (R1 and R2) with an average Phred quality score ≥30. R2 reads were reverse complemented, and both R1 and R2 were aligned independently to the C57BL/6 mouse kappa light chain V and J gene reference sequences from the IMGT^9^ database using IgBLAST^112^. The proportion of Vĸ gene usage was calculated. Statistical comparisons between genotypes were performed using the two-sided Mann–Whitney U test.

### Genome-wide RNA sequencing and analysis

Total RNA was extracted from ex vivo FACS-purified pre-B cells using the NucleoSpin RNA Kit according to the manufacturer’s instructions. RNA integrity was assessed using an Agilent TapeStation, and concentrations were quantified with a Qubit fluorometer (Thermo Fisher). A total of 1 µg of DNase-treated RNA was used. Ribosomal RNA was depleted prior to library construction. RNA-seq libraries were prepared using the SMARTer Stranded Total RNA Sample Prep Kit – HI Mammalian (Takara) following the manufacturer’s recommendations. This kit generates libraries compatible with Illumina platforms for mammalian samples. Libraries were pooled to a final concentration of 10 nM and loaded on a NextSeq 500 using the NextSeq 500/550 High-Output v2.5 Kit (75 cycles). Sequencing generated 122-bp single-end reads. Raw FASTQ files were quality-checked and trimmed using Trim Galore (v0.6.10). Trimmed reads were aligned to the mm10 reference genome using STAR (v2.7.11) with default parameters^113^. Resulting BAM files were used for gene-level quantification with HTSeq (v2.0.3)^114^. Raw read counts were analyzed in R (v4.4.1), and normalization and differential gene expression analysis were performed using the DESeq2 package^115^. Genes with a fold change >1.85 and a Benjamini– Hochberg–adjusted p-value (Padj) <0.05 were considered significantly differentially expressed.

### ATAC sequencing

ATAC-seq libraries were prepared from 50–100 × 10³ *ex vivo* FACS-purified pre-B cells using the Active Motif ATAC-Seq Kit (Active Motif, #53150) according to the manufacturer’s protocol. Briefly, nuclei were isolated, tagmented with pre-assembled transposomes, and DNA was purified. Libraries were amplified using the Nextera Illumina kit following the manufacturer’s instructions. Final libraries were size-selected with a 1.2× SPRI bead cleanup and sequenced on an Illumina NovaSeq 6000 platform with 61-bp paired-end reads.

### H3K27ac and CTCF ChIP sequencing

For H3K27ac ChIP-sequencing, approximately 1.8–5.3 × 10⁶ CD19⁺ pre-B cells were enriched from WT-Rag1⁻/⁻.hIgM and mutant-Rag1⁻/⁻.hIgM mice using anti-CD19 magnetic beads (Miltenyi Biotec) and MACS column separation (Miltenyi Biotec). For CTCF ChIP-sequencing, 3–6 × 10⁶ pre-B cells cultured in IL-7 from WT and ΔrsCBE mice were used. Cells were cross-linked with 1% formaldehyde for 10 min at room temperature and quenched with 0.1 M glycine. Cells were washed twice with cold phosphate-buffered saline (PBS) and lysed with lysis buffer [0.5% SDS, 10 mM EDTA, 50 mM Tris-HCl (pH 8), and protease inhibitor]. DNA was sonicated in an ultrasonic bath (Diagenode, Bioruptor) to an average length of 300–500 bp. Sonicated chromatin was centrifuged at 16,000 rpm for 15 min, and the supernatants were immunoprecipitated overnight with anti-H3K27ac (Active Motif, #39133) or anti-CTCF (Cell Signaling Technology, #3418S) antibodies. Protein G beads (Cell Signaling Technology, #9006) were added to immunoprecipitated samples and incubated for 2.5 h at 4°C. Beads were sequentially washed for 5 min each in low-salt RIPA [20 mM Tris-HCl (pH 8), 150 mM NaCl, 2 mM EDTA, 1% Triton X-100, 0.1% SDS], high-salt RIPA [20 mM Tris-HCl (pH 8), 500 mM NaCl, 2 mM EDTA, 1% Triton X-100, 0.1% SDS], LiCl buffer [10 mM Tris (pH 8.0), 1 mM EDTA, 250 mM LiCl, 1% NP-40, 1% Na-deoxycholate], and Tris-EDTA buffer. Beads were eluted in TE (pH 8.0) with 0.1% SDS and 150 mM NaCl for 1 h at 65 °C, followed by RNase treatment for 30 min at 37 °C. Proteinase K was added and incubated at 50 °C for 2 h. Reverse cross-linking was performed by adding NaCl to a final concentration of 200 mM and incubating for at least 6 h at 65°C. DNA was purified by phenol– chloroform extraction, and libraries were prepared from the immunoprecipitated DNA using the KAPA Hyper Prep Kit according to the manufacturer’s protocol. H3K27ac ChIP–seq libraries were sequenced on an Illumina NovaSeq 6000, generating 122-bp single-end reads, while CTCF ChIP–seq libraries were sequenced on an Illumina NextSeq, generating 122-bp single-end reads.

### ChIP-qPCR

To assess CTCF occupancy at the targeted deletion site, 445.3-WT and ΔrsCBE-18 bp 445.3 v-Abl Rag1-deficient cells were subjected to anti-CTCF ChIP analysis. Experiments were performed using 1 × 10⁷ cells per condition, following the protocols described above, with 1% formaldehyde. Immunoprecipitated and input DNA samples were analysed by quantitative real-time PCR, and enrichment was calculated relative to input to account for variability in chromatin quantity.

### ChIP sequencing and ATAC sequencing data analysis

Raw FASTQ files from ATAC-seq and ChIP-seq experiments (paired-end and single-end, respectively) were processed. Quality control and adapter trimming were performed using Trim Galore (v0.6.10). Cleaned reads were aligned to the mm10 reference genome using Bowtie2 (v2.5.2) with default parameters^116^, and the resulting alignments were saved as BAM files. Samtools (v1.19.2) merge was then used with default parameters to combine data from the two biological replicates^117^. The merged BAM files were converted to BigWig format for visualization using bamCoverage, normalized to counts per million (CPM)^118^. Peak calling was performed on the merged BAM files using MACS2 (v2.2.8) callpeak^119^, with peaks identified using MACS2 for ATAC-seq and de novo peak detection for ChIP-seq. Read counts from both experiments were normalized using the csaw package. The normalized counts were converted into a DGEList object in edgeR to identify differentially accessible or enriched regions^120^. Normalized BigWig signal intensities were extracted for defined genomic regions and visualized as heatmaps using plotHeatmap for ATAC-seq and H3K27ac ChIP–seq^118^.

### Micro-C-TALE sequencing

Micro-C-TALE was carried out as described below (unpublished data). Briefly, 1.6–4 × 10⁶ fixed and permeabilized cells were treated with micrococcal nuclease (MNase) (ThermoFisher, #88216) to obtain predominantly mononucleosomal DNA. The DNA was then ligated using T4 DNA ligase (NEB, #M0202L), extracted by phenol– chloroform and ethanol precipitation, and ligation products were enriched by size selection using Ampure XP beads (Beckman Coulter, #A63881). End repair and adaptor ligation were performed using the NEBNext Ultra II DNA Library Prep Kit (NEB, #E7645S), followed by PCR amplification with indexed primers. Four Micro-C libraries were generated and pooled in equimolar proportions.

Capture probes were prepared from fifteen BACs spanning the target genomic region. BAC DNA was mixed equimolarly, sheared to ∼200-bp fragments by sonication, end-repaired, and ligated to specialized adaptors using the NEBNext Ultra II kit. The BAC library was then PCR-amplified using biotinylated primers. Hybridization-based capture was performed by incubating the pooled Micro-C libraries with the biotinylated BAC-derived probes in multiple replicates. Hybridized products were pulled down using streptavidin-coated magnetic beads (Dynabeads MyOne sterptavidin C1, Invitrogen, #65001) and amplified by PCR with Illumina-compatible primers. The capture cycle was repeated once, and the final libraries were sequenced on a DNBSEQ-G400 using paired-end 150-bp reads.

### Micro-C-TALE analysis

Reads were mapped, filtered, and balanced using nf-distiller (https://github.com/open2c/distiller-nf) and cooler (https://academic.oup.com/bioinformatics/article/36/1/311/5530598).

### Apoptosis Assay

Apoptosis was assessed using the FITC Annexin V Apoptosis Detection Kit with propidium iodide (PI) (BioLegend, #640914). Fc receptors were blocked with anti-CD16/CD32 antibody in 1 × 10⁶ Histopaque-purified bone marrow lymphocytes. Following centrifugation, cells were stained with fluorochrome-conjugated antibodies against surface markers for 25 min at 4 °C, washed twice with FACS buffer, and resuspended in Annexin V binding buffer. Annexin V and PI were added (5 µl each per 100 µl of cell suspension) and incubated for 15 min at room temperature in the dark. Samples were then diluted with an additional 100 µl of binding buffer prior to flow cytometric acquisition.

### Cell cycle Analysis

Bone marrow cells were subjected to ACK lysis, followed by surface marker staining of 2 × 10⁶ cells with fluorochrome-conjugated antibodies for cell cycle analysis. Cells were then incubated with Hoechst 33342 (Sigma-Aldrich, #B2261) (1 µl of 5 mg ml⁻¹ stock per 500 µl DPBS) for 30 min at room temperature with mixing, resuspended in 200 µl FACS buffer, and analysed on a CytoFLEX flow cytometer.

### Serum isolation

Blood was collected from mice via tail vein or cardiac puncture into 1.5 ml microcentrifuge tubes. Samples were allowed to coagulate at room temperature (20– 25 °C) for 30–60 min, followed by centrifugation at 1,500 × g for 10 min. The resulting serum was carefully aspirated and stored at −80 °C until further analysis.

### ELISA

ELISA reactions were carried out using flat-bottom MaxiSorp™ 96-well Nunc-Immuno plates (Thermo Scientific, #442404). The assay was adapted from a previously published protocol^121^. Antigens (Hep-2 lysate; lipopolysaccharides from *Escherichia coli* O55:B5, Merck, #L4005-100MG; Salmon Sperm DNA Solution, Invitrogen, #15632011; Human Insulin Solution, Merck, #I9278-5ML) were coated in PBS at 100 µl per well and incubated overnight at 4 °C. For comparative ELISA assays with multiple targets, antigens were plated at 0.5 µg/well. Plates were washed five times with washing buffer (1× PBS, 0.05% Tween-20) and incubated with 100 µl blocking buffer (1× PBS with 1% BSA) for 1 h at room temperature. The blocking solution was then replaced with serum samples for 2 h at room temperature. Serum samples were assayed at a 1:50 dilution. Plates were washed five times with washing buffer and incubated with anti-mouse IgG secondary antibody conjugated to horseradish peroxidase (HRP) (Jackson ImmunoResearch, #115-035-062) in PBS at a 1:6,000 dilution. After an additional five washes, plates were developed using 3,3’,5,5’-tetramethylbenzidine (TMB) (BioFX TMB One Component HRP Microwell Substrate, Surmodics), and absorbance was measured at 630 nm using an ELISA plate reader (Tecan Spark).

### Quantification and Statistical analysis

Statistical analyses were performed using two-tailed unpaired t-tests in GraphPad Prism (v8.0.1). Data are expressed as mean ± SD, with p-values < 0.05 considered statistically significant. Details of specific statistical tests are provided in the corresponding sections.

## Acknowledgements

We thank Ihab Abd-Elrahman and Kirill Makedonski from the National Genetically Engineered Mouse Models (GEMM) Unit, The Hebrew University of Jerusalem, for generating the transgenic mice. We are grateful to A. Nasereddin and I. Shiff from the Genomic Applications Laboratory, Core Research Facility, Faculty of Medicine – Ein Kerem, Hebrew University of Jerusalem, for their scientific advice and RNA sequencing services. We also thank Dr. Hadas Segev-Yekutiel and Dr. Eleonora Medvedev from the Core Research Facility at the Hebrew University School of Medicine for their assistance with flow cytometry analysis. This work was supported by research grants from the Israel Academy of Sciences (grant no. 1228/18 to Y.B.), the Israel Cancer Research Foundation (grant no. 211410 to Y.B.), the Israel Cancer Association (grant no. 20241009 to Y.B.), the United States–Israel Binational Science Foundation (grant no. 2100289 to Y.B.), and the Emanuel Rubin Chair in Medical Sciences (Y.B.).

## Author contributions

D.G. conceived the study, designed and performed all experiments, and analysed and interpreted the data. E.G. performed bioinformatic analyses of all high-throughput datasets. D.G. and A.G. conducted the Micro-C-TALE sequencing experiments under the supervision of N.K. We adapted the Micro-C-TALE method from a protocol originally developed by AKG. D.G. performed the CTCF ChIP–seq and H3K27ac ChIP–seq experiments under the supervision of M.H and Y.D. A.H. provided the 4C sequencing scripts and primer information. Bar Avidov analysed the Vκ RNA repertoire for the rsCBE genotype and provided the corresponding analysis scripts. F.C. and L.H. contributed to the FACS experiments and offered valuable technical guidance for the analysis. Batia Azria contributed to the initial analysis of the RNA-seq dataset for the mutant genotypes. D.G. wrote the entire manuscript with input from all authors. Y.B. oversaw, supervised and directed the entire study.

## Data and materials availability

High-throughput sequencing data generated in this study have been deposited in the Gene Expression Omnibus (GEO) database. Igκ repertoire RNA-seq data are available under accession number GSE313431; genome-wide RNA-seq data under GSE313430; ATAC-seq data under GSE313428; CTCF ChIP-seq data under GSE313432; H3K27ac ChIP-seq data under GSE313433; 4C-seq data under GSE313429; and Micro-C-TALE-seq data under GSE313783.

## Extended Data Figures

**Extended Data Fig. 1:**
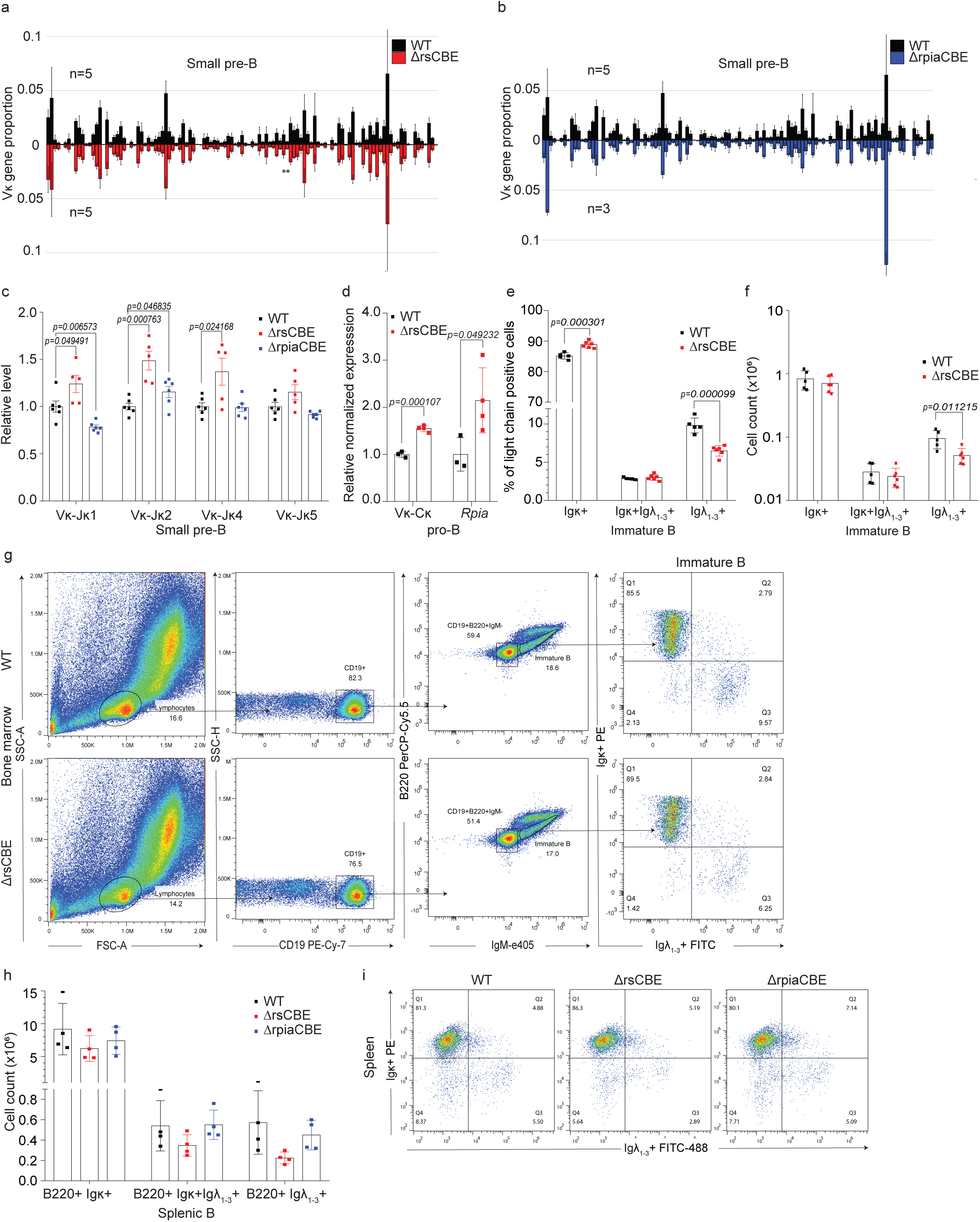
CBE is essential for maintaining light chain isotype balance and enabling bidirectional transcriptional insulation. **a, b,** Vκ gene rearrangement frequencies in RNA from bone marrow–derived small pre-B cells of ΔrsCBE and ΔrpiaCBE mice. Error bars represent s.d. for each Vκ gene. Statistical significance was determined by t-test (p-adj < 0.001, ***; p-adj < 0.01, **; p-adj < 0.05, *). **c,** Relative levels of rearranged Jκ segments in small pre-B cells from WT, ΔrsCBE and ΔrpiaCBE mice, measured by qPCR and normalized to the Eμ region (n = 5–6). **d,** *Rpia* and total Igκ transcript levels in bone marrow–derived pro-B cells from WT and ΔrsCBE mice, measured by qPCR and normalized to *Ubc* and *Ppia* (n = 3). **e,** Igκ and Igλ isotype exclusion in IgM⁺ B cells from WT and ΔrsCBE mice, shown as percentages; **f,** corresponding cell counts (in millions) (n = 5–6 mice per genotype). Absolute numbers of each B cell subset were quantified from the femur, tibia, and fibula of each mouse. **g,** Gating strategy for Igκ⁺ and Igλ⁺ Immature B cells in bone marrow from WT and ΔrsCBE mice; values shown as percentages in plots. **h,** Igκ and Igλ isotype exclusion in B220⁺ splenic B cells from WT, ΔrsCBE and ΔrpiaCBE mice, shown as cell counts (in millions) (n = 4). Data are mean ± s.d.; P values were calculated by unpaired Student’s t-test. Absolute numbers of each B cell subset were quantified from the femur, tibia, and fibula of each mouse. **i,** Representative gating strategy of B220⁺ splenic B cells for Igκ⁺ and Igλ⁺ populations, with percentages indicated in each quadrant.

**Extended Data Fig. 2:**
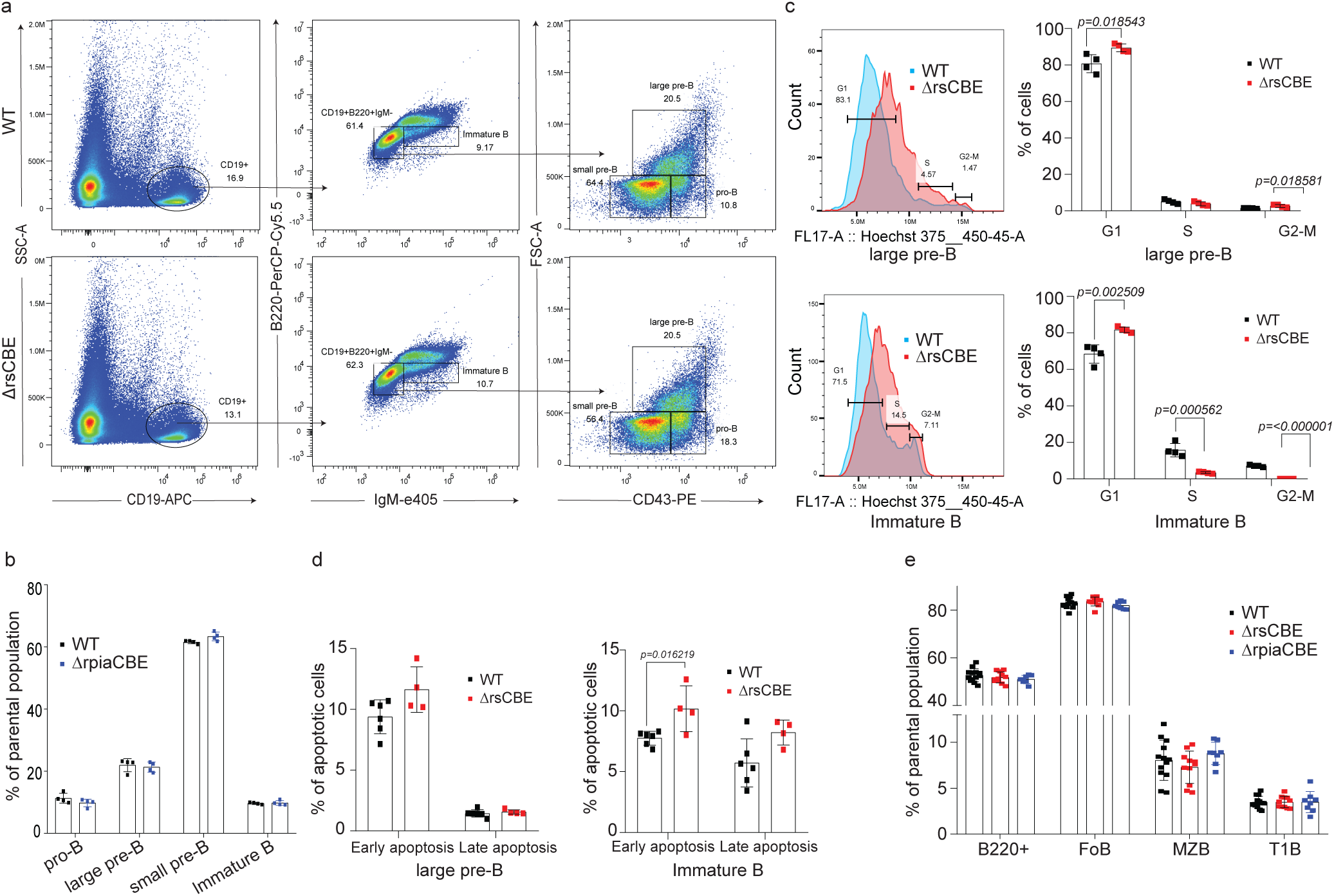
CBE controls early B cell development. **a,** Representative gating strategy for identifying bone marrow B cell subsets in WT and ΔrsCBE mice. **b,** Flow cytometric quantification of bone marrow B cell populations in WT and ΔrpiaCBE mice (n = 4), presented as percentages of the parental population. **c,** Hoechst-based cell-cycle analysis of bone marrow B cell subsets in WT and ΔrsCBE mice; percentages of cells in cycle (G1/S/G2–M DNA content) are shown (n = 4). **d,** Apoptosis in B cell subsets assessed by Annexin V and PI staining; early (Annexin V⁺) and late (Annexin V⁺ PI⁺) apoptotic cells are presented as percentages (n = 4). Data are mean ± s.d.; P values were determined using an unpaired t-test. **e,** Flow cytometric quantification of splenic B cell subsets in WT, ΔrsCBE and ΔrpiaCBE mice (n = 8–10), presented as percentages of the parental population (mean ± s.d.); P values were determined using an unpaired t-test.

**Extended Data Fig. 3:**
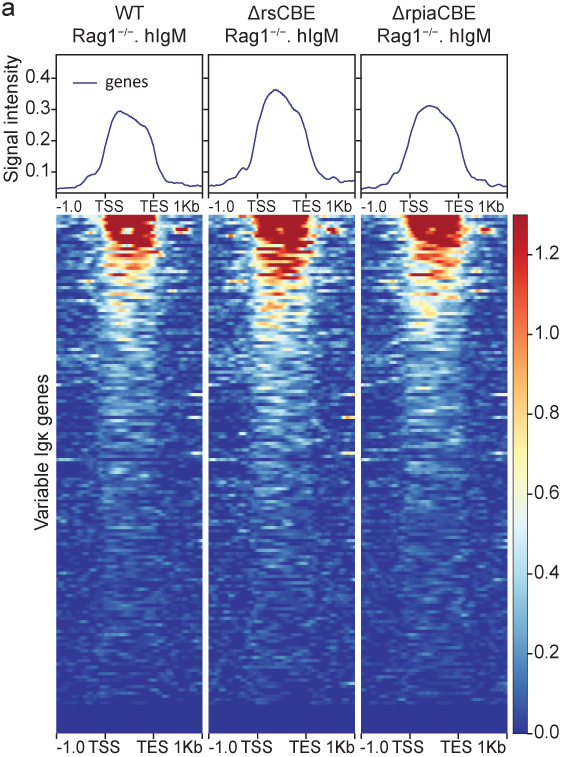
Co-ordinated molecular epigenetic mechanisms orchestrate Igκ expression. **a,** Heat map of H3K27ac enrichment across Vκ gene segments in WT-Rag1⁻/⁻.hIgM, ΔrsCBE-Rag1⁻/⁻.hIgM, and ΔrpiaCBE-Rag1⁻/⁻.hIgM pre-B cells. Signal intensity was quantified within a 2-kb window spanning 1 kb upstream of the transcription start site (TSS) to 1 kb downstream of the TES. Data are averages of two biological replicates.

**Extended Data Fig. 4:**
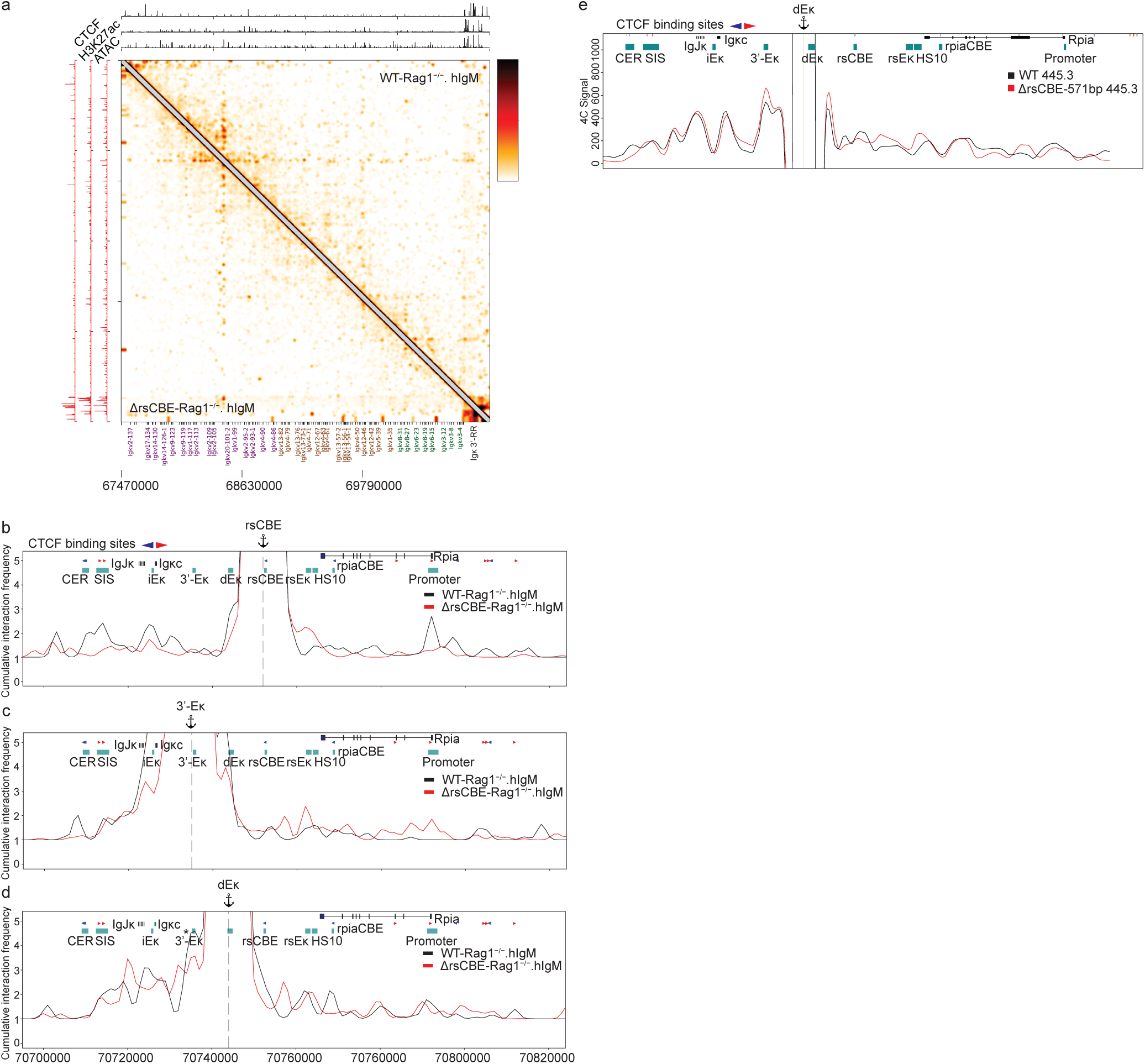
CBE controls the intricate Igκ super-enhancer architecture through chromatin insulation. **a,** Micro-C-TALE contact maps across the Igκ locus in WT-Rag1⁻/⁻.hIgM and ΔrsCBE-Rag1⁻/⁻.hIgM pre–B cells at 5-kb resolution and smoothed with a Gaussian filter (σ = 5 kb). **b,** Virtual 4C interaction profiles from Micro-C-TALE data (1-kb resolution, Gaussian smoothing, σ = 1 kb) in WT-Rag1⁻/⁻.hIgM and ΔrsCBE-Rag1⁻/⁻.hIgM pre–B cells with rsCBE **c**, 3′-Eκ, and **d,** dEκ as anchor viewpoints. **e,** 4C-seq interaction profiles across the 3′-RR with dEκ as the viewpoint in WT 445.3 and ΔrsCBE-571bp 445.3 cells. Interaction frequencies are shown as reads per million (RPM) with a 100-bp window and 21-bp running window. Data represent the mean of two biological replicates.

**Extended Data Fig. 5:**
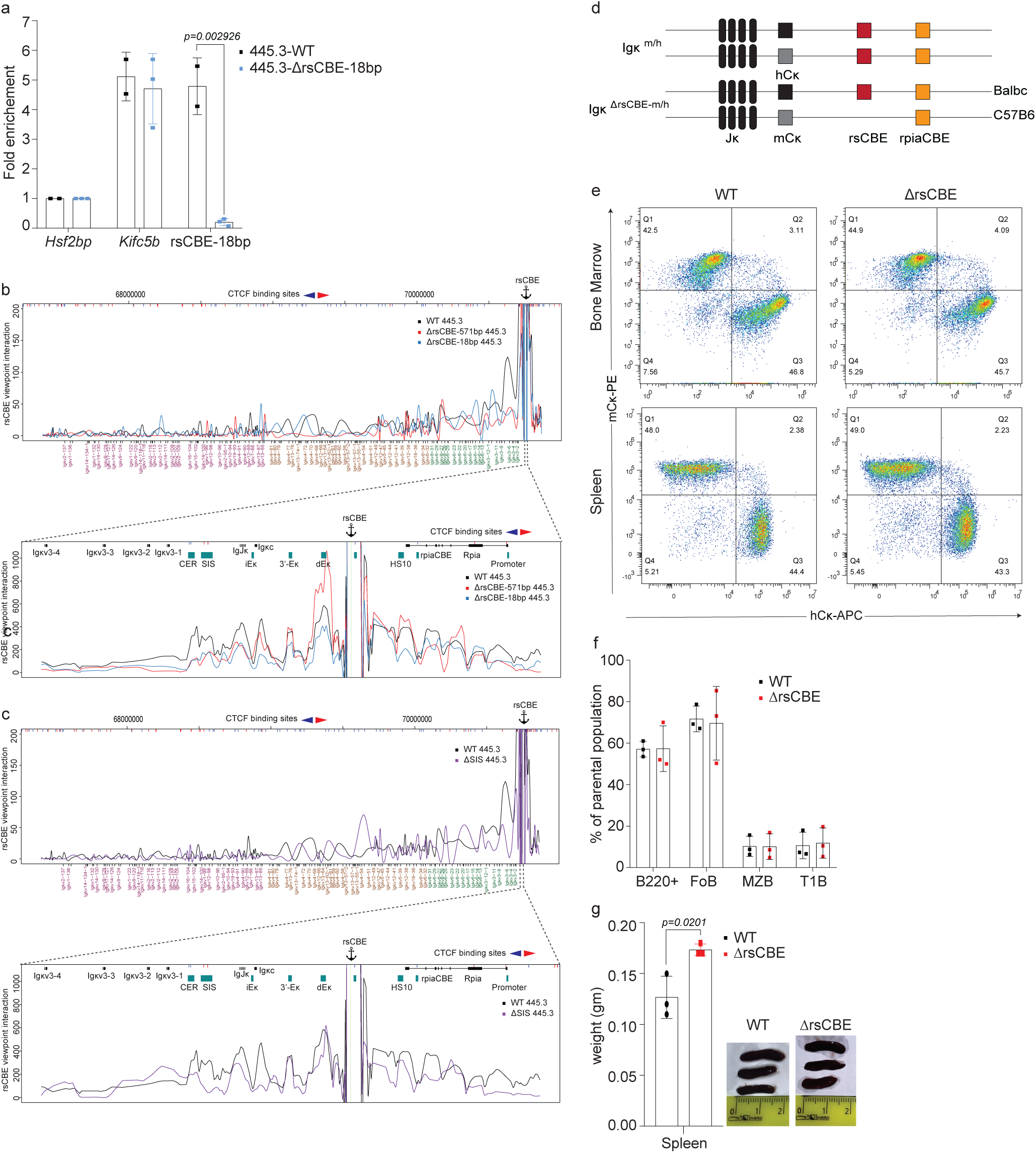
rsCBE-mediated chromatin loop extrusion governs B-cell tolerance and autoimmunity. **a,** CTCF occupancy at the rsCBE region in Rag1-deficient WT 445.3 and ΔrsCBE–18 bp 445.3 single-cell clones, measured by ChIP–qPCR and expressed as fold enrichment. The murine *Kifc5b* locus served as a positive control for CTCF occupancy, while *Hsf2bp* was used as a negative control lacking CTCF binding. Data are presented as mean ± s.d. (n = 2–3). Statistical significance was determined using an unpaired t-test; p < 0.05 was considered significant. **b,** 4C-seq interaction profiles across the Igκ locus in Rag1-deficient WT 445.3, ΔrsCBE-571 bp 445.3, ΔrsCBE-18 bp 445.3, and **c,** ΔSIS 445.3 cells. Interaction frequencies are shown as reads per million (RPM) using a 100-bp window with a 21-bp running window. The anchor denotes the rsCBE viewpoint, and a magnified view of the 3′-RR (dotted line) shows mean interaction profiles from two independent biological replicates. **d,** Schematic of mCκ⁺ and hCκ⁺ alleles in Igκ^m/h^ and Igκ^ΔrsCBE-m/h^ hybrid mice. **e,** Representative flow cytometric gating of bone marrow and splenic IgM⁺ B cells in hybrid mice, showing the proportions of mCκ⁺, hCκ⁺, and dual-mCκ⁺hCκ⁺ B cells within the gated populations. Values are shown as percentages in each quadrant. **f,** Flow cytometric quantification of splenic B cell populations in 14-month-old mice (n = 3), shown as percentages of the parental population. **g,** Splenic weight and images from 14-month-old mice (n = 3). P values were determined by unpaired t-test.

## References

1 Alt, F. W., Zhang, Y., Meng, F. L., Guo, C. & Schwer, B. Mechanisms of programmed DNA lesions and genomic instability in the immune system. Cell 152, 417–429, doi:10.1016/j.cell.2013.01.007 (2013).

2 Deguine, J. & Xavier, R. J. B cell tolerance and autoimmunity: Lessons from repertoires. J Exp Med 221, doi:10.1084/jem.20231314 (2024).

3 Young, C., Lau, A. W. Y. & Burnett, D. L. B cells in the balance: Offsetting self-reactivity avoidance with protection against foreign. Front Immunol 13, 951385, doi:10.3389/fimmu.2022.951385 (2022).

4 Watanabe, A. et al. Self-tolerance curtails the B cell repertoire to microbial epitopes. JCI Insight 4, doi:10.1172/jci.insight.122551 (2019).

5 Jung, D. & Alt, F. W. Unraveling V(D)J recombination; insights into gene regulation. Cell 116, 299–311, doi:10.1016/s0092-8674(04)00039-x (2004).

6 Schatz, D. G. & Swanson, P. C. V(D)J recombination: mechanisms of initiation. Annu Rev Genet 45, 167–202, doi:10.1146/annurev-genet-110410-132552 (2011).

7 Jhunjhunwala, S., van Zelm, M. C., Peak, M. M. & Murre, C. Chromatin architecture and the generation of antigen receptor diversity. Cell 138, 435–448, doi:10.1016/j.cell.2009.07.016 (2009).

8 Harrow, J. et al. GENCODE: producing a reference annotation for ENCODE. Genome Biol 7 **Suppl 1**, S4 1–9, doi:10.1186/gb-2006-7-s1-s4 (2006).

9 Lefranc, M. P. et al. IMGT, the international ImMunoGeneTics information system. Nucleic Acids Res 37, D1006–1012, doi:10.1093/nar/gkn838 (2009).

10 Brekke, K. M. & Garrard, W. T. Assembly and analysis of the mouse immunoglobulin kappa gene sequence. Immunogenetics 56, 490–505, doi:10.1007/s00251-004-0659-0 (2004).

11 Kleiman, E., Xu, J. & Feeney, A. J. Cutting Edge: Proper Orientation of CTCF Sites in Cer Is Required for Normal Jkappa-Distal and Jkappa-Proximal Vkappa Gene Usage. J Immunol 201, 1633–1638, doi:10.4049/jimmunol.1800785 (2018).

12 Aoki-Ota, M., Torkamani, A., Ota, T., Schork, N. & Nemazee, D. Skewed primary Igkappa repertoire and V-J joining in C57BL/6 mice: implications for recombination accessibility and receptor editing. J Immunol 188, 2305–2315, doi:10.4049/jimmunol.1103484 (2012).

13 Inlay, M., Alt, F. W., Baltimore, D. & Xu, Y. Essential roles of the kappa light chain intronic enhancer and 3’ enhancer in kappa rearrangement and demethylation. Nat Immunol 3, 463–468, doi:10.1038/ni790 (2002).

14 Zhou, X., Xiang, Y., Ding, X. & Garrard, W. T. A new hypersensitive site, HS10, and the enhancers, E3’ and Ed, differentially regulate Igkappa gene expression. J Immunol 188, 2722–2732, doi:10.4049/jimmunol.1102758 (2012).

15 Zhou, X., Xiang, Y., Ding, X. & Garrard, W. T. Loss of an Igkappa gene enhancer in mature B cells results in rapid gene silencing and partial reversible dedifferentiation. Mol Cell Biol 33, 2091–2101, doi:10.1128/MCB.01569-12 (2013).

16 Gorman, J. R. et al. The Ig(kappa) enhancer influences the ratio of Ig(kappa) versus Ig(lambda) B lymphocytes. Immunity 5, 241–252, doi:10.1016/s1074-7613(00)80319-2 (1996).

17 Levin-Klein, R., Kirillov, A., Rosenbluh, C., Cedar, H. & Bergman, Y. A novel pax5-binding regulatory element in the igkappa locus. Front Immunol 5, 240, doi:10.3389/fimmu.2014.00240 (2014).

18 Xu, Y., Davidson, L., Alt, F. W. & Baltimore, D. Deletion of the Ig kappa light chain intronic enhancer/matrix attachment region impairs but does not abolish V kappa J kappa rearrangement. Immunity 4, 377–385, doi:10.1016/s1074-7613(00)80251-4 (1996).

19 Barajas-Mora, E. M. & Feeney, A. J. Enhancers as regulators of antigen receptor loci three-dimensional chromatin structure. Transcription 11, 37–51, doi:10.1080/21541264.2019.1699383 (2020).

20 Ribeiro de Almeida, C., et al. The DNA-binding protein CTCF limits proximal Vkappa recombination and restricts kappa enhancer interactions to the immunoglobulin kappa light chain locus. Immunity 35, 501–513, doi:10.1016/j.immuni.2011.07.014 (2011).

21 Jain, S., Ba, Z., Zhang, Y., Dai, H. Q. & Alt, F. W. CTCF-Binding Elements Mediate Accessibility of RAG Substrates During Chromatin Scanning. Cell 174, 102–116 e114, doi:10.1016/j.cell.2018.04.035 (2018).

22 Xiang, Y., Zhou, X., Hewitt, S. L., Skok, J. A. & Garrard, W. T. A multifunctional element in the mouse Igkappa locus that specifies repertoire and Ig loci subnuclear location. J Immunol 186, 5356–5366, doi:10.4049/jimmunol.1003794 (2011).

23 Ebert, A., Hill, L. & Busslinger, M. Spatial Regulation of V-(D)J Recombination at Antigen Receptor Loci. Adv Immunol 128, 93–121, doi:10.1016/bs.ai.2015.07.006 (2015).

24 Beagan, J. A. & Phillips-Cremins, J. E. On the existence and functionality of topologically associating domains. Nat Genet 52, 8–16, doi:10.1038/s41588-019-0561-1 (2020).

25 Phillips-Cremins, J. E. et al. Architectural protein subclasses shape 3D organization of genomes during lineage commitment. Cell 153, 1281–1295, doi:10.1016/j.cell.2013.04.053 (2013).

26 Schwarzer, W. et al. Two independent modes of chromatin organization revealed by cohesin removal. Nature 551, 51–56, doi:10.1038/nature24281 (2017).

27 Fudenberg, G. et al. Formation of Chromosomal Domains by Loop Extrusion. Cell Rep 15, 2038–2049, doi:10.1016/j.celrep.2016.04.085 (2016).

28 Rao, S. S. P. et al. Cohesin Loss Eliminates All Loop Domains. Cell 171, 305–320 e324, doi:10.1016/j.cell.2017.09.026 (2017).

29 Sanborn, A. L. et al. Chromatin extrusion explains key features of loop and domain formation in wild-type and engineered genomes. Proc Natl Acad Sci U S A 112, E6456–6465, doi:10.1073/pnas.1518552112 (2015).

30 Rao, S. S. et al. A 3D map of the human genome at kilobase resolution reveals principles of chromatin looping. Cell 159, 1665–1680, doi:10.1016/j.cell.2014.11.021 (2014).

31 Vos, E. S. M. et al. Interplay between CTCF boundaries and a super enhancer controls cohesin extrusion trajectories and gene expression. Mol Cell 81, 3082–3095 e3086, doi:10.1016/j.molcel.2021.06.008 (2021).

32 Liu, N. Q. et al. WAPL maintains a cohesin loading cycle to preserve cell-type-specific distal gene regulation. Nat Genet 53, 100–109, doi:10.1038/s41588-020-00744-4 (2021).

33 Zhu, Y., Denholtz, M., Lu, H. & Murre, C. Calcium signaling instructs NIPBL recruitment at active enhancers and promoters via distinct mechanisms to reconstruct genome compartmentalization. Genes Dev 35, 65–81, doi:10.1101/gad.343475.120 (2021).

34 Denholtz, M. et al. Upon microbial challenge, human neutrophils undergo rapid changes in nuclear architecture and chromatin folding to orchestrate an immediate inflammatory gene program. Genes Dev 34, 149–165, doi:10.1101/gad.333708.119 (2020).

35 Lichtenstein, M., Keini, G., Cedar, H. & Bergman, Y. B cell-specific demethylation: a novel role for the intronic kappa chain enhancer sequence. Cell 76, 913–923, doi:10.1016/0092-8674(94)90365-4 (1994).

36 Levin-Klein, R. & Bergman, Y. Regulation of IgL Chain Recombination. Encyclopedia of Immunobiology, Vol 1: Development and Phylogeny of the Immune System, 71–77, doi:10.1016/B978-0-12-374279-7.01008-0 (2016).

37 Qian, J. et al. B cell super-enhancers and regulatory clusters recruit AID tumorigenic activity. Cell 159, 1524–1537, doi:10.1016/j.cell.2014.11.013 (2014).

38 Barajas-Mora, E. M. et al. A B-Cell-Specific Enhancer Orchestrates Nuclear Architecture to Generate a Diverse Antigen Receptor Repertoire. Mol Cell 73, 48–60 e45, doi:10.1016/j.molcel.2018.10.013 (2019).

39 Barajas-Mora, E. M. et al. Enhancer-instructed epigenetic landscape and chromatin compartmentalization dictate a primary antibody repertoire protective against specific bacterial pathogens. Nat Immunol 24, 320–336, doi:10.1038/s41590-022-01402-z (2023).

40 Zhang, Y. et al. Molecular basis for differential Igk versus Igh V(D)J joining mechanisms. Nature 630, 189–197, doi:10.1038/s41586-024-07477-y (2024).

41 Ribeiro de Almeida, C., Hendriks, R. W. & Stadhouders, R. Dynamic Control of Long-Range Genomic Interactions at the Immunoglobulin kappa Light-Chain Locus. Adv Immunol 128, 183–271, doi:10.1016/bs.ai.2015.07.004 (2015).

42 Xiang, Y., Park, S. K. & Garrard, W. T. Vkappa gene repertoire and locus contraction are specified by critical DNase I hypersensitive sites within the Vkappa-Jkappa intervening region. J Immunol 190, 1819–1826, doi:10.4049/jimmunol.1203127 (2013).

43 Liu, Z. et al. A recombination silencer that specifies heterochromatin positioning and ikaros association in the immunoglobulin kappa locus. Immunity 24, 405–415, doi:10.1016/j.immuni.2006.02.001 (2006).

44 Xiang, Y., Park, S. K. & Garrard, W. T. A major deletion in the Vkappa-Jkappa intervening region results in hyperelevated transcription of proximal Vkappa genes and a severely restricted repertoire. J Immunol 193, 3746–3754, doi:10.4049/jimmunol.1401574 (2014).

45 Matheson, L. S. et al. Local Chromatin Features Including PU.1 and IKAROS Binding and H3K4 Methylation Shape the Repertoire of Immunoglobulin Kappa Genes Chosen for V(D)J Recombination. Front Immunol 8, 1550, doi:10.3389/fimmu.2017.01550 (2017).

46 Kleiman, E., Loguercio, S. & Feeney, A. J. Epigenetic Enhancer Marks and Transcription Factor Binding Influence Vkappa Gene Rearrangement in Pre-B Cells and Pro-B Cells. Front Immunol 9, 2074, doi:10.3389/fimmu.2018.02074 (2018).

47 Stadhouders, R. et al. Pre-B cell receptor signaling induces immunoglobulin kappa locus accessibility by functional redistribution of enhancer-mediated chromatin interactions. PLoS Biol 12, e1001791, doi:10.1371/journal.pbio.1001791 (2014).

48 Lazorchak, A. S., Schlissel, M. S. & Zhuang, Y. E2A and IRF-4/Pip promote chromatin modification and transcription of the immunoglobulin kappa locus in pre-B cells. Mol Cell Biol 26, 810–821, doi:10.1128/MCB.26.3.810-821.2006 (2006).

49 Vela, J. L., Ait-Azzouzene, D., Duong, B. H., Ota, T. & Nemazee, D. Rearrangement of mouse immunoglobulin kappa deleting element recombining sequence promotes immune tolerance and lambda B cell production. Immunity 28, 161–170, doi:10.1016/j.immuni.2007.12.011 (2008).

50 Durdik, J., Moore, M. W. & Selsing, E. Novel kappa light-chain gene rearrangements in mouse lambda light chain-producing B lymphocytes. Nature 307, 749–752, doi:10.1038/307749a0 (1984).

51 Martin, D. J. & van Ness, B. G. Initiation and processing of two kappa immunoglobulin germ line transcripts in mouse B cells. Mol Cell Biol 10, 1950–1958, doi:10.1128/mcb.10.5.1950-1958.1990 (1990).

52 Amin, R. H. et al. Biallelic, ubiquitous transcription from the distal germline Igkappa locus promoter during B cell development. Proc Natl Acad Sci U S A 106, 522–527, doi:10.1073/pnas.0808895106 (2009).

53 Nieh, Y. C. et al. Suppression of Ribose-5-Phosphate Isomerase a Induces ROS to Activate Autophagy, Apoptosis, and Cellular Senescence in Lung Cancer. Int J Mol Sci 23, doi:10.3390/ijms23147883 (2022).

54 Heintze, J., Costa, J. R., Weber, M. & Ketteler, R. Ribose 5-phosphate isomerase inhibits LC3 processing and basal autophagy. Cell Signal 28, 1380–1388, doi:10.1016/j.cellsig.2016.06.015 (2016).

55 Ciou, S. C. et al. Ribose-5-phosphate isomerase A regulates hepatocarcinogenesis via PP2A and ERK signaling. Int J Cancer 137, 104–115, doi:10.1002/ijc.29361 (2015).

56 Chou, Y. T. et al. Identification of a noncanonical function for ribose-5-phosphate isomerase A promotes colorectal cancer formation by stabilizing and activating beta-catenin via a novel C-terminal domain. PLoS Biol 16, e2003714, doi:10.1371/journal.pbio.2003714 (2018).

57 Novobrantseva, T. I. et al. Rearrangement and expression of immunoglobulin light chain genes can precede heavy chain expression during normal B cell development in mice. J Exp Med 189, 75–88, doi:10.1084/jem.189.1.75 (1999).

58 Espinoza, C. R. & Feeney, A. J. Chromatin accessibility and epigenetic modifications differ between frequently and infrequently rearranging VH genes. Mol Immunol 44, 2675–2685, doi:10.1016/j.molimm.2006.12.002 (2007).

59 Pulivarthy, S. R. et al. Regulated large-scale nucleosome density patterns and precise nucleosome positioning correlate with V(D)J recombination. Proc Natl Acad Sci U S A 113, E6427–E6436, doi:10.1073/pnas.1605543113 (2016).

60 Yancopoulos, G. D., Blackwell, T. K., Suh, H., Hood, L. & Alt, F. W. Introduced T cell receptor variable region gene segments recombine in pre-B cells: evidence that B and T cells use a common recombinase. Cell 44, 251–259, doi:10.1016/0092-8674(86)90759-2 (1986).

61 Yancopoulos, G. D. & Alt, F. W. Developmentally controlled and tissue-specific expression of unrearranged VH gene segments. Cell. 1985. 40: 271–281. *J Immunol* **188**, 10-20 (2012).

62 Stanhope-Baker, P., Hudson, K. M., Shaffer, A. L., Constantinescu, A. & Schlissel, M. S. Cell type-specific chromatin structure determines the targeting of V(D)J recombinase activity in vitro. Cell 85, 887–897, doi:10.1016/s0092-8674(00)81272-6 (1996).

63 McBlane, F. & Boyes, J. Stimulation of V(D)J recombination by histone acetylation. Curr Biol 10, 483–486, doi:10.1016/s0960-9822(00)00449-8 (2000).

64 Nightingale, K. P. et al. Acetylation increases access of remodelling complexes to their nucleosome targets to enhance initiation of V(D)J recombination. Nucleic Acids Res 35, 6311–6321, doi:10.1093/nar/gkm650 (2007).

65 Goldmit, M. et al. Epigenetic ontogeny of the Igk locus during B cell development. Nat Immunol 6, 198–203, doi:10.1038/ni1154 (2005).

66 Lin, Y. C. et al. Global changes in the nuclear positioning of genes and intra- and interdomain genomic interactions that orchestrate B cell fate. Nat Immunol 13, 1196–1204, doi:10.1038/ni.2432 (2012).

67 Crane, E. et al. Condensin-driven remodelling of X chromosome topology during dosage compensation. Nature 523, 240–244, doi:10.1038/nature14450 (2015).

68 Hill, L. et al. Igh and Igk loci use different folding principles for V gene recombination due to distinct chromosomal architectures of pro-B and pre-B cells. Nat Commun 14, 2316, doi:10.1038/s41467-023-37994-9 (2023).

69 Muljo, S. A. & Schlissel, M. S. A small molecule Abl kinase inhibitor induces differentiation of Abelson virus-transformed pre-B cell lines. Nat Immunol 4, 31–37, doi:10.1038/ni870 (2003).

70 Tiegs, S. L., Russell, D. M. & Nemazee, D. Receptor editing in self-reactive bone marrow B cells. J Exp Med 177, 1009–1020, doi:10.1084/jem.177.4.1009 (1993).

71 Retter, M. W. & Nemazee, D. Receptor editing occurs frequently during normal B cell development. J Exp Med 188, 1231–1238, doi:10.1084/jem.188.7.1231 (1998).

72 Moore, M. W., Durdik, J., Persiani, D. M. & Selsing, E. Deletions of kappa chain constant region genes in mouse lambda chain-producing B cells involve intrachromosomal DNA recombinations similar to V-J joining. Proc Natl Acad Sci U S A 82, 6211–6215, doi:10.1073/pnas.82.18.6211 (1985).

73 Dunda, O. & Corcos, D. Recombining sequence recombination in normal kappa-chain-expressing B cells. J Immunol 159, 4362–4366 (1997).

74 Zou, Y. R., Takeda, S. & Rajewsky, K. Gene targeting in the Ig kappa locus: efficient generation of lambda chain-expressing B cells, independent of gene rearrangements in Ig kappa. EMBO J 12, 811–820, doi:10.1002/j.1460-2075.1993.tb05721.x (1993).

75 Daitch, L. E., Moore, M. W., Persiani, D. M., Durdik, J. M. & Selsing, E. Transcription and recombination of the murine RS element. J Immunol 149, 832–840 (1992).

76 Nadel, B., Cazenave, P. A. & Sanchez, P. Murine lambda gene rearrangements: the stochastic model prevails over the ordered model. EMBO J 9, 435–440, doi:10.1002/j.1460-2075.1990.tb08128.x (1990).

77 Dorner, T., Foster, S. J., Farner, N. L. & Lipsky, P. E. Immunoglobulin kappa chain receptor editing in systemic lupus erythematosus. J Clin Invest 102, 688–694, doi:10.1172/JCI3113 (1998).

78 Casellas, R. et al. Igkappa allelic inclusion is a consequence of receptor editing. J Exp Med 204, 153–160, doi:10.1084/jem.20061918 (2007).

79 Liu, S. et al. Receptor editing can lead to allelic inclusion and development of B cells that retain antibodies reacting with high avidity autoantigens. J Immunol 175, 5067–5076, doi:10.4049/jimmunol.175.8.5067 (2005).

80 Gerdes, T. & Wabl, M. Autoreactivity and allelic inclusion in a B cell nuclear transfer mouse. Nat Immunol 5, 1282–1287, doi:10.1038/ni1133 (2004).

81 Velez, M. G. et al. Ig allotypic inclusion does not prevent B cell development or response. J Immunol 179, 1049–1057, doi:10.4049/jimmunol.179.2.1049 (2007).

82 Fournier, E. M. et al. Dual-reactive B cells are autoreactive and highly enriched in the plasmablast and memory B cell subsets of autoimmune mice. J Exp Med 209, 1797–1812, doi:10.1084/jem.20120332 (2012).

83 Panigrahi, A. K. et al. RS rearrangement frequency as a marker of receptor editing in lupus and type 1 diabetes. J Exp Med 205, 2985–2994, doi:10.1084/jem.20082053 (2008).

84 Fraser, L. D. et al. Immunoglobulin light chain allelic inclusion in systemic lupus erythematosus. Eur J Immunol 45, 2409–2419, doi:10.1002/eji.201545599 (2015).

85 Lipsky, P. E. Systemic lupus erythematosus: an autoimmune disease of B cell hyperactivity. Nat Immunol 2, 764–766, doi:10.1038/ni0901-764 (2001).

86 Grammer, A. C. & Lipsky, P. E. B cell abnormalities in systemic lupus erythematosus. Arthritis Res Ther 5 **Suppl 4**, S22–27, doi:10.1186/ar1009 (2003).

87 Kitaura, Y. et al. Control of the B cell-intrinsic tolerance programs by ubiquitin ligases Cbl and Cbl-b. Immunity 26, 567–578, doi:10.1016/j.immuni.2007.03.015 (2007).

88 Fritzler, M. J. Clinical relevance of autoantibodies in systemic rheumatic diseases. Mol Biol Rep 23, 133–145, doi:10.1007/BF00351161 (1996).

89 Takeda, S. et al. Deletion of the immunoglobulin kappa chain intron enhancer abolishes kappa chain gene rearrangement in cis but not lambda chain gene rearrangement in trans. EMBO J 12, 2329–2336, doi:10.1002/j.1460-2075.1993.tb05887.x (1993).

90 Chen, J. et al. B cell development in mice that lack one or both immunoglobulin kappa light chain genes. EMBO J 12, 821–830, doi:10.1002/j.1460-2075.1993.tb05722.x (1993).

91 Wong, J. B. et al. B-1a cells acquire their unique characteristics by bypassing the pre-BCR selection stage. Nat Commun 10, 4768, doi:10.1038/s41467-019-12824-z (2019).

92 Hu, J. et al. Chromosomal Loop Domains Direct the Recombination of Antigen Receptor Genes. Cell 163, 947–959, doi:10.1016/j.cell.2015.10.016 (2015).

93 Ba, Z. et al. CTCF orchestrates long-range cohesin-driven V(D)J recombinational scanning. Nature 586, 305–310, doi:10.1038/s41586-020-2578-0 (2020).

94 Lin, S. G., Ba, Z., Alt, F. W. & Zhang, Y. RAG Chromatin Scanning During V(D)J Recombination and Chromatin Loop Extrusion are Related Processes. Adv Immunol 139, 93–135, doi:10.1016/bs.ai.2018.07.001 (2018).

95 Zhao, L. et al. Orientation-specific RAG activity in chromosomal loop domains contributes to Tcrd V(D)J recombination during T cell development. J Exp Med 213, 1921–1936, doi:10.1084/jem.20160670 (2016).

96 Zhang, Y. et al. The fundamental role of chromatin loop extrusion in physiological V(D)J recombination. Nature 573, 600–604, doi:10.1038/s41586-019-1547-y (2019).

97 Dai, H. Q. et al. Loop extrusion mediates physiological Igh locus contraction for RAG scanning. Nature 590, 338–343, doi:10.1038/s41586-020-03121-7 (2021).

98 Zhang, Y., Zhang, X., Dai, H. Q., Hu, H. & Alt, F. W. The role of chromatin loop extrusion in antibody diversification. Nat Rev Immunol 22, 550–566, doi:10.1038/s41577-022-00679-3 (2022).

99 Hill, L. et al. Wapl repression by Pax5 promotes V gene recombination by Igh loop extrusion. Nature 584, 142–147, doi:10.1038/s41586-020-2454-y (2020).

100 Morel, L. et al. Functional dissection of systemic lupus erythematosus using congenic mouse strains. J Immunol 158, 6019–6028 (1997).

101 Spanopoulou, E. et al. Functional immunoglobulin transgenes guide ordered B-cell differentiation in Rag-1-deficient mice. Genes Dev 8, 1030–1042, doi:10.1101/gad.8.9.1030 (1994).

102 Xu, H., Li, H., Suri-Payer, E., Hardy, R. R. & Weigert, M. Regulation of anti-DNA B cells in recombination-activating gene-deficient mice. J Exp Med 188, 1247–1254, doi:10.1084/jem.188.7.1247 (1998).

103 Halverson, R., Torres, R. M. & Pelanda, R. Receptor editing is the main mechanism of B cell tolerance toward membrane antigens. Nat Immunol 5, 645–650, doi:10.1038/ni1076 (2004).

104 Hartley, S. B. et al. Elimination of self-reactive B lymphocytes proceeds in two stages: arrested development and cell death. Cell 72, 325–335, doi:10.1016/0092-8674(93)90111-3 (1993).

105 Lang, J. et al. Enforced Bcl-2 expression inhibits antigen-mediated clonal elimination of peripheral B cells in an antigen dose-dependent manner and promotes receptor editing in autoreactive, immature B cells. J Exp Med 186, 1513–1522, doi:10.1084/jem.186.9.1513 (1997).

106 Strasser, A. et al. Enforced BCL2 expression in B-lymphoid cells prolongs antibody responses and elicits autoimmune disease. Proc Natl Acad Sci U S A 88, 8661–8665, doi:10.1073/pnas.88.19.8661 (1991).

107 Ait-Azzouzene, D. et al. An immunoglobulin C kappa-reactive single chain antibody fusion protein induces tolerance through receptor editing in a normal polyclonal immune system. J Exp Med 201, 817–828, doi:10.1084/jem.20041854 (2005).

108 Nussenzweig, M. C. et al. Allelic exclusion in transgenic mice that express the membrane form of immunoglobulin mu. Science 236, 816–819, doi:10.1126/science.3107126 (1987).

109 Rolink, A., Kudo, A., Karasuyama, H., Kikuchi, Y. & Melchers, F. Long-term proliferating early pre B cell lines and clones with the potential to develop to surface Ig-positive, mitogen reactive B cells in vitro and in vivo. EMBO J 10, 327–336, doi:10.1002/j.1460-2075.1991.tb07953.x (1991).

110 Krijger, P. H. L., Geeven, G., Bianchi, V., Hilvering, C. R. E. & de Laat, W. 4C-seq from beginning to end: A detailed protocol for sample preparation and data analysis. Methods 170, 17–32, doi:10.1016/j.ymeth.2019.07.014 (2020).

111 Levin-Klein, R. et al. Clonally stable Vkappa allelic choice instructs Igkappa repertoire. Nat Commun 8, 15575, doi:10.1038/ncomms15575 (2017).

112 Ye, J., Ma, N., Madden, T. L. & Ostell, J. M. IgBLAST: an immunoglobulin variable domain sequence analysis tool. Nucleic Acids Res 41, W34–40, doi:10.1093/nar/gkt382 (2013).

113 Dobin, A. et al. STAR: ultrafast universal RNA-seq aligner. Bioinformatics 29, 15–21, doi:10.1093/bioinformatics/bts635 (2013).

114 Anders, S., Pyl, P. T. & Huber, W. HTSeq--a Python framework to work with high-throughput sequencing data. Bioinformatics 31, 166–169, doi:10.1093/bioinformatics/btu638 (2015).

115 Love, M. I., Huber, W. & Anders, S. Moderated estimation of fold change and dispersion for RNA-seq data with DESeq2. Genome Biol 15, 550, doi:10.1186/s13059-014-0550-8 (2014).

116 Langmead, B. & Salzberg, S. L. Fast gapped-read alignment with Bowtie 2. Nat Methods 9, 357–359, doi:10.1038/nmeth.1923 (2012).

117 Danecek, P. et al. Twelve years of SAMtools and BCFtools. Gigascience 10, doi:10.1093/gigascience/giab008 (2021).

118 Ramirez, F. et al. deepTools2: a next generation web server for deep-sequencing data analysis. Nucleic Acids Res 44, W160–165, doi:10.1093/nar/gkw257 (2016).

119 Zhang, Y. et al. Model-based analysis of ChIP-Seq (MACS). Genome Biol 9, R137, doi:10.1186/gb-2008-9-9-r137 (2008).

120 Reske, J. J., Wilson, M. R. & Chandler, R. L. ATAC-seq normalization method can significantly affect differential accessibility analysis and interpretation. Epigenetics Chromatin 13, 22, doi:10.1186/s13072-020-00342-y (2020).

121 Mazor, R. D. et al. Tumor-reactive antibodies evolve from non-binding and autoreactive precursors. Cell 185, 1208–1222 e1221, doi:10.1016/j.cell.2022.02.012 (2022).

